# Convergent evolution of noxious heat sensing by TRPA5, a novel class of heat sensor in *Rhodnius prolixus*

**DOI:** 10.1101/2023.05.26.542450

**Authors:** Marjorie A. Liénard, David Baez-Nieto, Cheng-Chia Tsai, Wendy A. Valencia-Montoya, Balder Werin, Urban Johanson, Jean-Marc Lassance, Jen Q. Pan, Nanfang Yu, Naomi E. Pierce

## Abstract

As ectotherms, insects need a multifaceted repertoire of heat-sensitive receptors to monitor environmental temperatures and finely control behavioral thermoregulation. Here, we show that *TRPA5* genes, a class of ankyrin transient receptor potential channels lost in genomes of model fruit flies or mosquitoes, are widespread across insect orders, and encode a previously uncharacterized type of heat receptors. We demonstrate that RpTRPA5B, a TRPA5 channel of the triatomine bug *Rhodnius prolixus* (Insect: Hemiptera), primary vector of Chagas disease, forms a homo-tetrameric channel displaying a uniquely high thermosensitivity. The channel biophysical determinants include a large channel activation enthalpy change (72 kcal/mol), a high temperature coefficient (Q_10_ = 25), and temperature-induced currents from 53 °C to 68 °C (T_0.5_= 58.6 °C) *in vitro,* similar to mammalian noxious TRPV heat receptors. Monomeric and tetrameric predictions of the ion channel architecture show reliable and conserved structural parallels with fruit fly dTRPA1, albeit depicting structural uniqueness from dTRPA, Painless and Pyrexia in the ankyrin repeat domain and the channel selectivity filter, potential modulator regions of functional characteristics of TRPs. The channel activation response, structural features and ubiquitous sensory tissue expression delineate a potential thermosensitive physiological niche close to that of *Pyrexia* genes, lost during the evolution of true bugs. Overall, the finding of *TRPA5* genes as a class of temperature-activated receptor illustrates the dynamic evolution of a large family of insect molecular heat detectors, with TRPs as promising multimodal sensory targets for triatomine vector control.

## Introduction

Animal thermosensation is critical for performance in fluctuating environments. Changes in environmental temperature are transduced by the sensory system as part of physiological feedback controlling responses such as metabolic homeostasis, feeding, finding suitable habitats, and extreme-heat avoidance (1, 2). At the molecular level, thermal perception is mediated by the temperature-dependent activation of specific cold- and heat-activated receptors (3, 4). Although families such as ionotropic receptors (IRs) and gustatory receptors (GRs) have been linked to peripheral innocuous thermosensation in insects (3–6), the transient receptor potential (TRP) receptor family encodes the greatest diversity of thermosensitive channels. TRP receptors are remarkably diverse (TRPA, TRPC, TRPN, TRPM, TRPML, and TRPV) and play salient roles as polymodal ion channels responding to chemical, mechanical, and thermal stimuli (7–12).

Mammalian TRP channels involved in temperature detection (thermoTRPs) belong to the TRPA, TRPV and TRPM subfamilies and are activated by temperatures from noxious cold to noxious heat (e.g., 4, 9, 13-16) (Table 1 and references therein). In invertebrates, known thermoTRP channels have so far been restricted to the ankyrin TRPA subfamily of genes, including *Painless*, *Pyrexia*, *TRPA1*, and *HsTRPA* (Hymenoptera-specific) (Fig. 1A; Fig. S1) (11, 12). In *Drosophila melanogaster, Painless*, *Pyrexia* and *dTRPA1* isoforms A, B and D encode receptors that exhibit distinct biophysical properties, cellular expression patterns, and temperature activation thresholds ranging from 19°C to 46°C (17–24). TRPA1 is also a heat–activated TRP sensor in *Anopheles gambiae* (25-37°C), and other mosquitos (25, 26), playing a key role in tuning heat-seeking behaviour. Outside the Diptera, TRPA1 has been characterized as a heat-sensitive channel in other insects as it is known to regulate the induction of embryonic diapause in *Bombyx mori* at temperatures above 21°C (27). The subfamily *Waterwitch* (*Wtrw*) includes receptors responding to stimuli in different modalities, from ancestral hygrosensation found in fruit flies (20) to derived heat sensing exhibited by hymenopterans and mediated by the HsTRPA subfamily, which diverged following a duplication from *Wtrw* (12). Thus, despite the loss of TRPA1 in Hymenoptera, in the honeybee, *Apis mellifera* Am-HsTRPA responds to temperatures around 34°C, and in the fire ant *Solenopsis invicta,* Si-HsTRPA is activated in the range 28-37 °C, whereas in the parasitoid wasp *Nasonia vitripennis,* Nv-HsTRPA activates in response to small temperature differences in the range 8 °C to 44 °C regardless of initial temperatures (28, 29). Notably, the insect TRP ankyrin family has an additional subfamily of unknown function, TRPA5, which is absent from the fruit fly genome yet found across several other orders of insects (11) (Fig. 1A).

**Figure 1.**
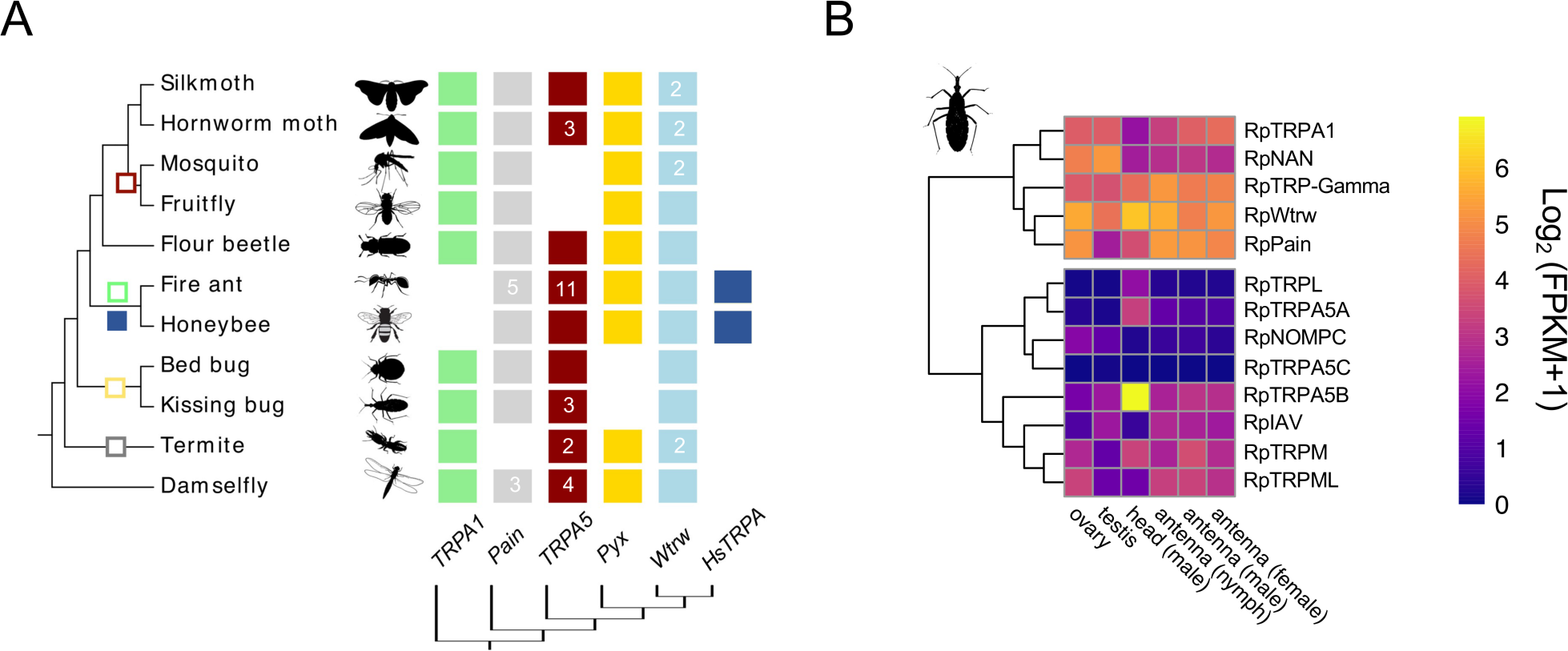
A. Phylogenetic reconstruction of the ankyrin TRP (TRPA) channel subfamilies in representative insect species. TRPA5 channels are present across insect Orders but absent from dipteran genomes (see also Figs. S1 and S2). Gene abbreviations: *Painless (Pain), Pyrexia (Pyx), Waterwitch (Wtrw*), TRPA Hymenoptera-specific (*HsTRPA*). Silkmoth, *Bombyx mori*; Hornworm moth, *Manduca sexta*; Mosquito, *Anopheles gambiae*; Fruit fly, *Drosophila melanogaster*; Flour beetle, *Tribolium castaneum*; Fire ant, *Solenopsis invicta*; Honeybee, *Apis mellifera*; Bed bug, *Cimex lectularis;* Kissing bug*, Rhodnius prolixus*; Termite, *Zootermopsis nevadensis*; Bluetail Damselfly, *Ischnura elegans.* Gene gain, filled square; gene loss, empty square. Numbers within squares indicate gene number when different from 1. **B.** TRP genes in *R. prolixus* and their relative expression levels across tissues in compiled transcriptomic data (*see Methods*). Heat maps compare the expression levels across tissues and developmental stages. Expression levels are represented as Log_2_ FPKM +1 and depicted with a gradient color scale. Gene models are based on genomic annotations (36), and *de novo* transcriptome assembly (e.g., 43) (see *Methods,* Table S2).

**Table 1.**
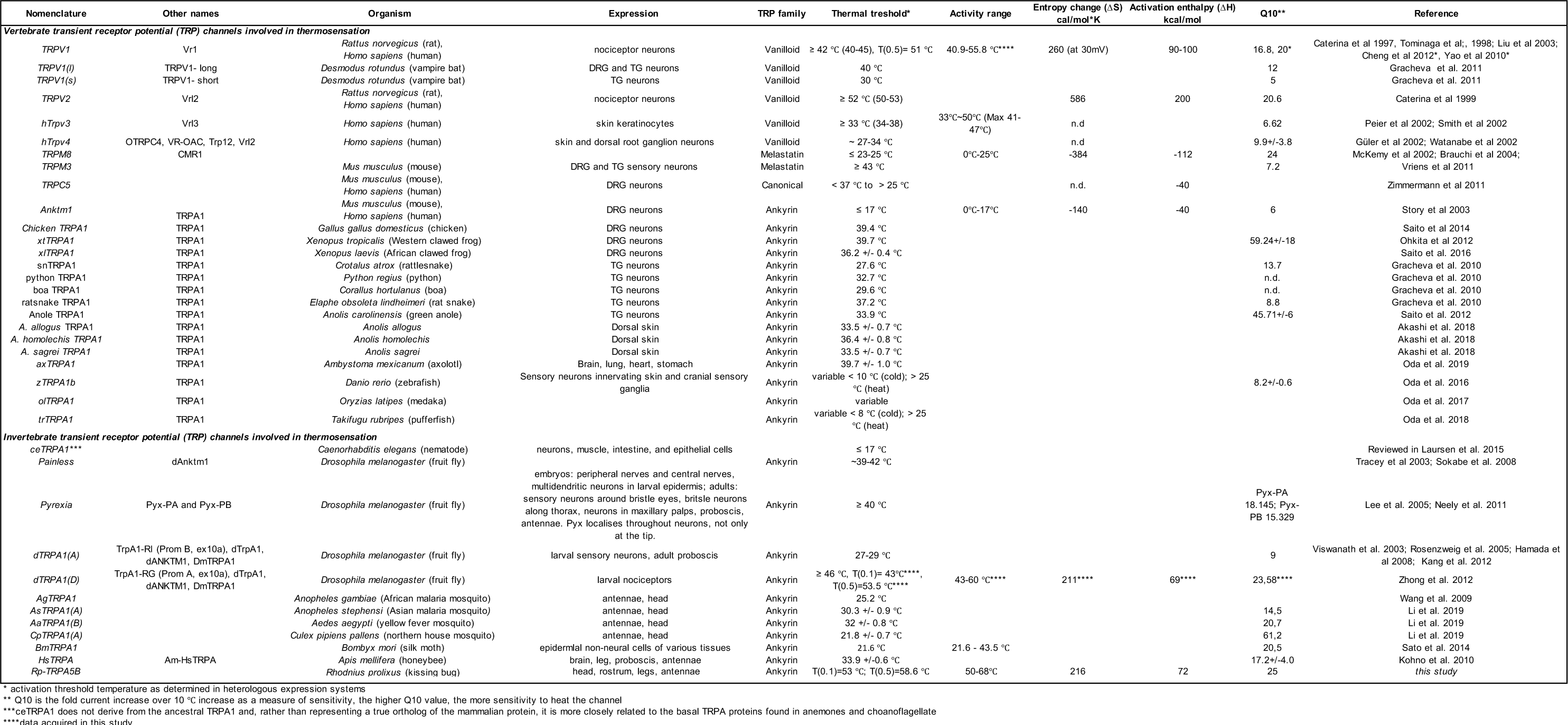
Vertebrate and invertebrate TRP ion channels involved in thermal transduction.

Here, we de-orphanize and characterize an ankyrin TRPA5 ion channel from the triatomine bug, *Rhodnius prolixus*. Long used as a model organism in studies of insect development and physiology (30), *R. prolixus* (Hemiptera; Reduviidae: Triatominae) has become increasingly relevant for molecular and functional studies. This is primarily explained by its long-term medical and societal impact as a haematophagous vector of *Trypanosoma cruzi*, the causative agent of Chagas’ disease (31). Due to the progressive adaptation of wild triatomine vector species to domestic environments, vector transmission to human populations has increased in recent years (32, 33). The disease currently affects over 8 million people worldwide, with vector transmission causing around 30,000 new cases and 12,000 deaths per year (33–35). Extensive long-term efforts towards decoding the sensory ecology of triatomines (30, 36, 37) have identified olfactory, thermal and environmentally-mediated cues as well as the neuroethology underlying its complex host-seeking behaviour (37–44). Moreover, the annotated *R. prolixus* genome (36) and recent transcriptomic studies (43–45) provide detailed profiles of candidate sensory receptor genes, including olfactory, ionotropic, pickpocket, and transient receptor potential receptors that can be used to probe the genetic basis of sensory traits (46, 47).

By leveraging genomic and transcriptomic resources available for *R. prolixus* along with molecular, structural modelling and functional approaches, we characterize a broadly expressed TRPA5 ion channel. The biophysical properties of the ion channel demonstrate that *RpTRPA5B* encodes an ankyrin type of heat-activated TRP receptor responding to noxious temperatures *in vitro*. Analyses of predicted structures reveal that the channel displays shared conserved structural domains with other ankyrin TRPs combined with unique features among the ankyrin family. Our findings open new avenues for future studies to assist in the development of multimodal tools for triatomine vector control efforts.

## Results

### Genomic and phylogenetic placement of Rhodnius ankyrin TRPs

To begin investigating the molecular basis of thermosensation in *Rhodnius prolixus*, we reanalysed the genome annotation (Version RproC3.3) complemented with transcriptomic resources (*see Methods*) to gain insights into gene variation and genomic architecture within the *R. prolixus* TRP ankyrin family. Genomes of triatomines (36) and additional surveyed hemipteran species (Fig. 1A, Fig. S1, Table S1) harbour members of four TRPA subfamilies. All surveyed species appear to lack an ortholog to *Pyrexia* (*Pyx*) TRP but possess one gene copy of three canonical ankyrin TRP genes: *Waterwitch* (*Wtrw*), *TRPA1* and *Painless* (*Pain*) (Fig. 1A, Fig. S1). Three *TRPA5* transcripts were previously described in *R. prolixus* (43). We updated the genome annotation of the reference assembly using transcriptomic datasets and show that *TRPA5A* (RPRC001596) and *TRPA5B* (RPRC001597) map to different genomic locations on a single scaffold and consist of two physically close tandem-duplicate loci, whereas *TRPA5C* (RPRC000570) maps to a distinct scaffold. Intrigued by the finding of multiple *TRPA5* gene copies, we performed an extensive *TRPA5* gene search across annotated genomic and transcriptomic datasets available for the insect Orders Anoplura, Diptera, Coleoptera, Hemiptera, Hymenoptera, Isoptera, Lepidoptera, Odonata and Thysanoptera. Our phylogenetic reconstruction shows that the TRPA5 ankyrin subfamily is completely absent in all surveyed dipteran genomes (Fig. S2), but TRPA5 orthologues are present at least in the orders Lepidoptera, Coleoptera, Hymenoptera, Hemiptera, Isoptera and Odonata (Fig. 1A, Fig. S2).

### Transcriptomic and quantitative expression of *TRPA5*

We next analyzed RNA-Seq raw data to assess the expression profile of *TRPAs* for *Rhodnius prolixus* (Fig. 1B). *TRPA1, Waterwitch* and *Painless* appear broadly and highly expressed. The three *TRPA5* genes differed more in their expression pattern: RpTRPA5A and RpTRPA5C mRNAs are expressed at low detection thresholds, and RpTRPA5B mRNA is moderately abundant across the range of surveyed tissues, including male head and adult antennae (Fig. 1B). Intrigued by high head expression and using complementary analyses by quantitative PCR, we confirmed that RpTRPA5B is also highly expressed in female heads, and ubiquitously expressed in the thorax, abdomen, rostrum, and legs (Fig. 1B, Fig. S3), which directed our choice toward this TRP gene for functional analyses on this channel subfamily.

### Validation of a functional assay using whole-cell patch clamp and optical heat-pulse delivery

In order to demonstrate the potential role of candidate TRPA5B as a thermosensitive ion channel, we transiently expressed a bicistronic T2A-fluorescent marker cassette (48) together with the candidate TRP channel, which localized well to the plasma membrane (Fig. 2A; Figs. S4) and optimized an *in vitro* cell-based workflow to record temperature-elicited currents from HEK293T cells under whole-cell patch-clamp configuration (Fig. 2B-C). Fast temperature stimulus was delivered by coupling an infrared laser diode to fiber optic after Yao et al. (49). A proportional-integral-derivative (PID) controller was used to keep the temperature stable along the duration of the pulse (700 ms), turning on and off the diode using the open pipette current trace as feedback for the PID controller, and calculating the steady-state parameters of activation from the current at the end of the 700 ms temperature pulse. The magnitude of ionic current changes through the open patch-clamp pipette was used to calculate the temperature changes associated with the different laser intensities (Fig. 2B, Fig. S5, Fig. S6). During this calibration, the laser voltage input and the series of pulses necessary to reach the desired temperatures are recorded, and this file is later played back to the diode. The patch-clamp recording pipette is positioned in the exact same position relative to the optic fiber during the calibration (Fig. S6), and each cell recorded has its own calibration file (see *Methods*). To validate this modified infrared (IR) patch-clamp system and expression cassette, we first transiently expressed two control thermoTRPs, the rat TRPV1 (rTRPV1) and fruit fly TRPA1 isoform D (dTRPA1-D) (Fig 2D-F, Fig. S5). At the molecular level, both rTRPV1 and dTRPA1-D formed expected homotetrameric structures (Fig. S4).

**Figure 2.**
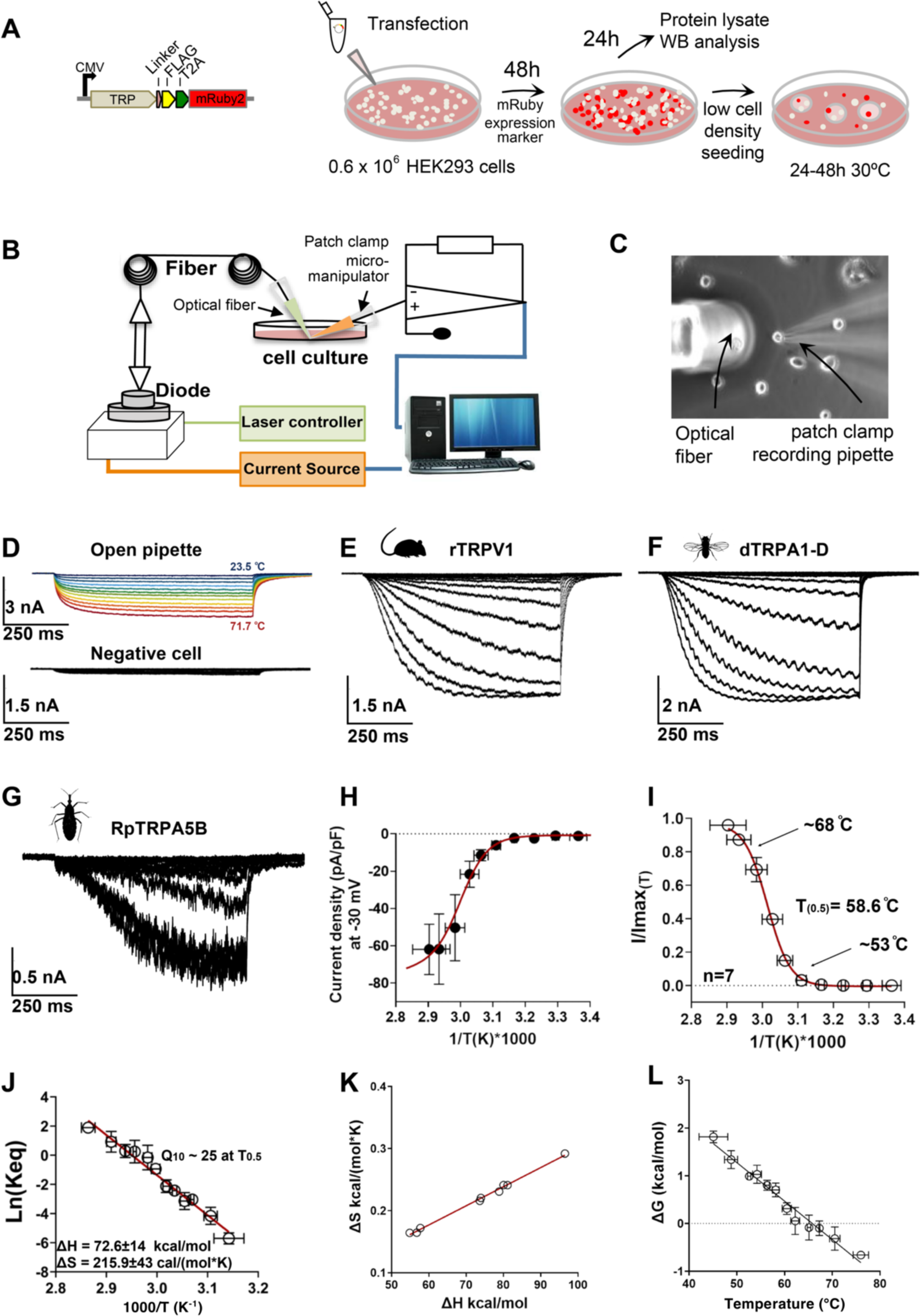
Thermodynamics of RpTRPA5B temperature-activated currents. A-C. Experimental workflow. **A.** Each TRP channel subcloned in the pFRT-TO-FLAG-T2A-mRuby2 expression cassette (48, 50) was transfected in HEK293T cells seeded at low density and incubated at 37°C for 48h. Cells were then prepared for patch-clamp recording by seeding in a 30-mm^2^ culture dish overlaid with round glass cover slips and incubated at 30 °C. **B.** Electrophysiology recordings took place after 24h to 48h using an optical fiber-based setup adapted after Yao et al 2010 (49), designed to couple manual patch clamp recordings with fiber optics as a way to provide controllable optical and thermal stimulations to individual cells expressing candidate thermosensitive receptor proteins. The setup consists of a fiber launch system combining a high-power optical fiber tuned to near-infrared wavelengths (λc =1460 nm (+/-20 nm), Po= 4.8 watts), a visible alignment laser (red), and a laser diode controller, forming a PID control loop using the patch clamp current as the feedback signal. **C**. During the experiment, a laser spot is aligned with one single patched cell (see Fig. S6) stably expressing the membrane receptor protein of interest in the cover slip placed in the recording chamber. **D.** *Upper panel*, current traces through the open patch-clamp pipette in response to temperature calibration steps from room temperature up to 71 °C elicited by increments in the IR laser voltage input (see *Methods*). Each 700 ms voltage pulse is represented in different colors for the different temperatures calculated from the open pipette currents. *Lower panel*, representative recording of non-transfected cells, these cells did not show robust temperature-elicited currents, like negative cells on the recording plate **E.** Whole-cell currents evoked by temperature steps from HEK293T cells expressing rat TRPV1 (heat-activated mammalian vanilloid thermoTRP), cells were held at -30 mV during the recording. **F.** Whole-cell currents evoked by temperature steps from HEK293T cells expressing dTRPA1-D (holding potential of -30 mV). The sinusoidal pattern observed within the current curves is inherent to the cyclic modulation of the laser’s rapid ‘on-off’ cycles. **G**. Whole-cell currents evoked by temperature steps in HEK293T cells expressing RpTRPA5B, cells were held at -30 mV. **H**. Current-Temperature relationship for RpTRPA5B whole-cell current was normalized by cell membrane capacitance (current density), the red line corresponds to a modified Boltzmann function that includes the leak and unitary current temperature dependence (see *Methods*). **I.** Fraction of RpTRPA5B channels in the open state (open probability, P_o_) as a function of the temperature. The Po vs 1/T was fitted to a Boltzman function with the midpoint of activation (T_0.5_) reached at 58.6 °C. **J.** van’t Hoff plot estimates of RpTRPA5B with an activation enthalpy of the endothermic transition at 92 kcal/mol and an entropic change associated with the temperature activation process at 274 cal/mol*K at -30 mV (49). **K**. Coupling between enthalpic (ΔH) and entropic (ΔS) changes for each one of the experiments recorded **L**. Free energy (ΔG) associated with the activation process as a function of temperature for RpTRP5AB channels. The receptor activation is associated with small free energy changes, as reported before for other families of mammalian thermoTRP receptors. ΔG was calculated as -RT*ln(Keq) (51). Data is represented as mean ± standard error.

### Full current activation response profiles at high temperatures for rTRPV1 and dTRPA1-D

Current pulses were set to result in temperature increments at the cell membrane in the range of 23.5 °C to 71.7 °C, at a holding membrane potential of -30 mV (Fig. 2D). This voltage magnitude provides a driving force big enough to resolve the ionic currents and minimizes a potential influence of the membrane voltage over the temperature activation process (49). A similar laser stimulation protocol led to marginal whole-cell current changes in non-transfected cells (Fig. 2D, Fig. S5A-B). Compared to non-transfected cells, we then observed a strong increase in the current amplitude of cells expressing rTRPV1 (Fig. 2E, Fig. S5C-D) with an enthalpy change associated with the activation of 88.3 ± 9.4 kcal/mol, which is comparable to the published enthalpy values obtained using millisecond temperature jumps of ΔH = 85 kcal/mol for rTPRV1 (49) (see Table 1) (13, 49, 52). The temperature values as shown in Fig. S5 for rTRPV1 align with those reported in earlier studies for this channel. The published threshold of activation for rTRPV1 is in the range 40-42 °C and corresponds to the temperature at which the first observable currents were detected (13, 53, 54). Hence, the channel emerging temperature recorded in our setup is at 40.9 °C when the first activation currents emerge over the dotted line (Fig. S5D). We calculated values of rTRPV1 T0.1 (i.e., the temperature for which there is a probability for 10% of channels to be open) at 45.3 °C (-30mV), and T=0.5 at 51.6°C (-30mV), consistently with values reported in Yao et al 2010 (49) (T0.5 51°C, -60mV). A temperature-induced activation response was also observed for the heat-activated fruit fly channel, dTRPA1-D, for which in our more precise setup, at 46.3 °C (22), the open probability (Po) of the channel is about 10% (Po=0.1), corroborating a higher-than-ambient activation temperature > 42 °C (22). Assuming complete activation of this channel (Po=1) by temperature, which was not measured in previous studies due to limitations in the maximum temperature to which the dTRPA1-D channel could be subjected, the activation process is characterized by an enthalpy change ΔH= 68.7 ± 13.1 kcal/mol and T_0.5_= 53.5 °C (Fig. 2F, Fig. S5E-F).

### Controlled temperature-dependent biophysical properties of RpTRPA5B

RpTRPA5B similarly assembled as a membrane-bound homotetramer when expressed in HEK293T cells (Fig. S4). When holding the membrane potential at -30mV in patched mRuby2-expressing cells transfected with RpTRPA5B, whole-cell currents were evoked by temperature steps a little above 50 °C (Fig. 2G, 2I). The average temperature for the activation “threshold” was 53°C, defined as Po = 0.1 calculated from the van’t Hoff plots. The channel opening appeared to saturate at 68°C (Po=0.9) (Fig. 2I), with a T_0.5_= 58.6°C. The current density versus temperature relationship (Fig. 2H) indicates that the opening of RpTRPA5B involves an activation enthalpy (ΔH) of approximately 72.6 ± 14 kcal/mol (Fig. 2J). The large entropy value (Fig. 2J, 2K) further indicates that the channel transits between a highly ordered closed state and a strongly disordered open configuration, close to activation enthalpy for TRPV1 (ΔH 101 ± 4 kcal/mol at -60mV) (49). Based on van’t Hoff plots thermodynamic parameters, we further calculated a Q_10_ value of ∼ 25, which is in the range of characterized noxious vertebrate receptors (rTRPV1 Q_10_ = 16.8; rTRPV2 Q_10_ = 20.6) and the invertebrate fruit fly Pyrexia (Q_10_ = 18.2) (*see also* Table 1).

### Insights from Rhodnius TRPA5B monomeric and tetrameric structure predictions

To visualize and compare structural features between TRPA5B and other ankyrin TRP homologues, we used AlphaFold 2.0 to generate models without a structural template (55–57). This approach was first reliably validated by a comparison to the recently reported structure of *Drosophila melanogaster* dTRPA1-A in state-1, which confirmed all distinct predicted features in the monomeric model of dTRPA1, including the interfacial helix and the interaction between ankyrin repeat (AR) 12 and the region C-terminal of the coiled-coil helix (Fig. 3A) (58). Although some of the helices in the monomeric model are oriented in unrealistic directions owing to missing constraints of the other monomers and the interactions that would force the C-terminus into the coiled-coil, their reliable secondary structure provides a first meaningful comparison of secondary structural elements and the general fold between proteins from different ankyrin TRP subfamilies.

**Figure 3.**
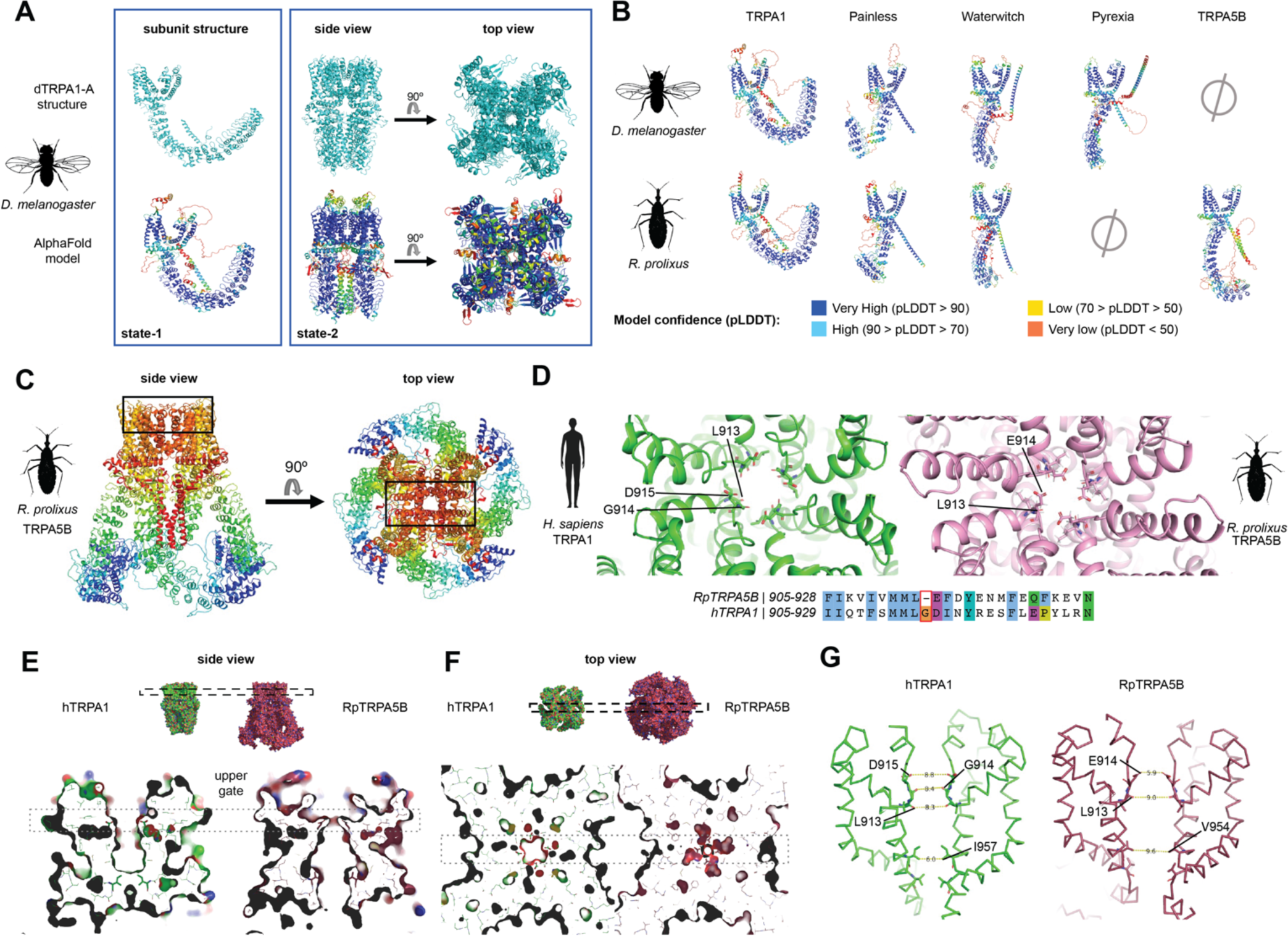
**Monomeric and tetrameric assemblies of RpTRPA5B channels modelled using AlphaFold, after validation with dTRPA1 structure. A**. *(left panel) Upper row,* cartoon representation of chain A in tetramer of dTRPA1 in state 1 (PDB ID 7YKR). The fold of a monomer in the experimentally determined structure of the dTRPA1 tetramer is very similar to the fold of an AlphaFold model of a single dTRPA1 monomer (*bottom row*). AlphaFold model colored from red to blue according to pLDDT confidence scores as shown in (B). The low-confidence regions (red) are not resolved in the reported structure and are likely to be intrinsically disordered. (*Right panel) Upper row,* experimentally determined structure of dTRPA1 in state 2 (PDB ID 7YKS). *Bottom row*, tetrameric AlphaFold model of dTRPA1 depicted as cartoons colored from red to blue according to confidence as in (B). The N- and C-terminal regions, which are not resolved in 7YKS, were excluded in the prediction. Only the last five of the 17 ankyrin repeats (AR12-16) are visible in the structure and overall regions with low confidence in the model (red-yellow) are not resolved in the structure. **B.** Monomers of *Drosophila* and *Rhodnius* TRPAs coloured by pLDDT score from the AlphaFold modelling. **C.** Tetrameric model of RpTRPA5B, coloured as chain bows (N-terminus, blue; C-terminus, red). The black box indicates the location of the pore and selectivity filter shown in **(D)**. **D.** Top view of the selectivity filter of the pore of hTRPA1 (*left*: human TRPA1, PDB ID 6V9Y) and RpTRPA5B (*right*). Three important residues(60) – L913, G914 and D915 – are marked in hTRPA1. The equivalent residues L913 and E914 are marked in RpTRPA5B, and G914 absent in RpTRPA5B is highlighted in the sequence alignment, together with additional residue changes adjacent to the selectivity filter. **E-F.** Comparison of the pore in hTRPA1 and model of RpTRPA5B indicates a closed upper gate and an open lower gate in the RpTRPA5B model. **E.** *(upper row)* Surface representation of hTRPA1 (green; PDB ID 6V9Y) and RpTRPA5B (pink), side view. The dashed box indicates the location of the upper gate towards the outside of the cell. *(lower row)* Slab along the pore through the transmembrane domain of the hTRPA1 structure and RpTRPA5B model. **F.** *(upper row)* Top view of hTRPA1 and RpTRPA5B shown in **E**. *(lower row)* The slab is perpendicular to the pore at the level of the upper gate shown in **E**. The dashed box indicates the location of slab in **(E). G**. Distances between corresponding residues in the upper and lower gate in structures of hTRPA1 and the model of RpTRPA5B, shown as sticks.

All *Rhodnius* and *Drosophila* ankyrin TRP monomeric structures were then modelled following the same approach, supporting highly reliable predictions for orthologues (Fig. 3B, Fig. S7) and expected structural similarities with the cryo-EM structures of *Drosophila* dTRPA1 and human hTRPA1(59), including the N-terminal ankyrin repeat (AR) domain, six transmembrane α-helices (S1-S6), and a region corresponding to the allosteric nexus of hTRPA1 connecting the AR domain and the transmembrane region (59). In addition, the monomeric N-termini show an overall conserved stacking of the ankyrin repeats (Fig. 3B, Fig. S7) - *albeit* with clade-specific breaks and numbers of repeats (Table S5). The C-terminal regions feature at least one α-helix, which together with the corresponding helices from the other subunits, most likely form a coiled-coil in the tetramers as seen in the solved TRPA1 structures.

We next generated a tetrameric model of RpTRPA5B to examine the predicted stable state of the pore and selectivity filter (Fig. 3D-G), following supporting evidence that an AlphaFold tetrameric model of dTRPA1 generated without a structural template proved highly comparable to the corresponding resolved structure of dTRPA1 in state-2 including the transmembrane domain, last part of the ARD (ARD12-16) and coiled-coil regions (Fig. 3A, see Fig. S8). For this, the root mean square deviation of the 20 Cα atoms of residues in the upper and lower gate of the pore was low (0.328 Å) (Fig. S8D, Table S6) with the sole main deviation found in the side chain of Glu982 (Fig. S9D). However, in the selectivity filter in the upper gate of the RpTRPA5B tetramer model (Fig. 3C and D), one glycine suggested to be important in gating (60) (Gly914 in hTRPA1, PDB:6V9Y), is absent in RpTRPA5B (Fig 3D), while conserved in most other TRPAs or substituted for serine or threonine in most other non-hemipteran TRPA5 proteins (see Fig. S2). Other adjacent residue changes are located in this region, and while Leu913 and Glu914 in RpTRPA5B form a shorter pore loop, they maintain the same overall locations with Leu913 and Asp915 in hTRPA1. Finally, an interesting last feature is the pore of the RpTRPA5B model, which appears to be open at the lower gate and closed at the upper gate (Fig. 3E-G, Fig. S9, Table S6), reversely to the hTRPA1 and dTRPA1structures (6V9Y, 7YKS).

## Discussion

### *TRPA5* evolutionary dynamics and function support insect thermoTRP channel usage plasticity

Large scale phylogenetic reconstructions combining TRP genes from 46 insect families spanning 9 major Orders provided additional insights into the dynamic evolution of five insect TRPA ankyrin clades (11, 12), *TRPA1*, *Painless*, *Waterwitch* (including *HsTRPA1*), *Pyrexia,* and *TRPA5* (Fig 1, Fig. S1, Table S1). In addition to complex alternative splicing (61), remarkable group-specific expansions of *TRPA5* genes as observed in the fire ant, *Solenopsis invicta* (29), in the damselfly *Ischnura elegans*, the tobacco hornworm moth *Manduca sexta*, and several hemipteran species including the kissing bug, *Rhodnius prolixus*, seem to play a role in the TRPA5 clade diversification (Fig. 1, Figs S1-S2). Whereas the TRPA5 clade is absent in all surveyed mosquitoes and flies (Diptera), reversely, *Pyrexia* genes, encoding a class of functional noxious heat receptors in fruit flies (18), are retained in all major insects except in hemipterans. Dynamic gene loss-gain amongst insect TRPA lineages together with experimental evidence of a hemipteran thermosensitive TRPA5 support channel usage plasticity and convergence in noxious thermal activation range over millions of years of divergent evolution. Altogether, our findings illustrate an example of the resilience of invertebrate sensory systems via compensatory molecular sensors of environmental thermal detection.

In *Rhodnius*, *TRPA* expression (Fig. 1B) matches the tissue distribution range of insect thermoreceptors such as the canonical fruit fly *TRPA1*, *Painless* and *Pyrexia* (27–29, 62). For instance, *Painless* is expressed along the entire epidermis in fruit fly larvae (17) and *Pyrexia* is expressed in sensory structures more broadly, including adult mouth structures (maxillary palps and proboscis), adult bristle sensilla, thorax, and eyes, and larval body epidermis (18). Different dTRPA1 isoforms localize in different tissues, including brain neurons and blood capillaries (isoform A) (21, 63) larval nociceptive neurons in the central nervous system (brain, isoform D) (22), and proboscis (isoform C) (23). Fruit fly *Wtrw* (a humidity-sensor) and mosquito *TRPA1* localize to specific antennal sensilla (20, 25), and honeybee and fire ant *HsTRPA1* orthologues are both expressed broadly including in leg, antenna, head, and proboscis (honeybee) (28), or antenna, leg, head, thorax and abdomen (fire ant) (29). Although all *Rhodnius TRPA* genes likely have physiological roles, including *RpTRPA5A*, and *RpTRPA5C*, we chose to focus first on *TRPA5B* as representative to start investigating the biophysical properties of TRPA5 channels, guided by transcriptomic and quantitative tissue expression analyses that placed *TRPA5B* as an interesting broadly-expressed TRP found across the adult body of *Rhodnius* (Fig. 1B, Fig. S3), including tissues known to have potential thermosensing roles.

### TRPA5B is gated by noxious temperature stimuli *in vitro*

By delivering controlled optical heat-pulses to HEK293T cells expressing TRP proteins under whole cell voltage-clamp configuration at -30mV, we first recapitulated reliably the biophysical properties associated to thermal activation of two control thermoTRP channels, the rat rTRPV1 and the fruit fly dTRPA1-D (Fig 2, Fig. S5). For the dTRPA1-D channel, our setup permitted extended biophysical characterization to obtain T_0.5_ = 53.5 °C (ΔH = 69 kcal/mol, -30 mV), a widely accepted comparative measure of the temperature at which is achieved the probability of having 50% channels open, not calculated in earlier studies due to setup constraints (22). For rTRPV1, our data consistently recapitulated previously reported activation thresholds above 40 °C (40.9 °C, Fig. S5D) (13, 53) and a value of T_0.5_ = 51.6 °C (-30 mV) (Fig. S5D) (49, 54), establishing a rigorous basis for temperature stimulus delivery to determine the biophysical properties of RpTRPA5B thermal activation. For RpTRPA5B, whole-cell currents were evoked above 50 °C with the probability of having 10 % channels open (Po = 0.1) at 53 °C, a T_0.5_= 58.6 °C, and saturating currents were reached at 68 °C, representing Po=0.9 (Fig. 2). High Q_10_ values for this channel at T_0.5_ were 25, comparable to values of Q_10_ 16-20 for rTRPV1 (13, 49, 53, 54, 64) and Q_10_ 20.6 for rTRPV2 (15), Q_10_ 15-18 for fruit fly Pyx (18, 62), and Q_10_ 23 for dTRPA1 (Fig. 2, Table 1) whereas non-thermo-TRP channels typically have Q_10_ values below 3 (9), supporting that RpTRPA5B is gated by temperature stimuli.

The relatively large observed enthalpy change (ΔH = 72.6 ± 14 kcal/mol at -30 mV) (Fig. 3) required for the channel activation is also in line with a high sensitivity to temperature changes. Hence, high enthalpy changes of 100 kcal/mol and 88kcal/mol are required to activate rTRPV1 at -60 mV (49, 54) and at -30 mV (Fig. S5D), respectively, representing a three-fold increase in temperature sensitivity at negative voltages (49) compared to its activation in depolarizing conditions (30 kcal/mol at +60 mV). Similarly, RpTRPA5B showed a robust response at negative voltages (-30 mV) and almost no heat-elicited activity at depolarized potentials (> 0 mV), supporting its dependence on temperature for activation. Large enthalpy changes ranging from 60 to 200 kcal/mol are also involved in the opening of other highly temperature-dependent channels including dTRPA1-D (Fig. S5F), TRPM8, TRPV1, and TRPV2 (65).

The maximum Po value for TRPA5B’s thermal activation (Fig. 3) is lower compared to the dTRPA1-D and rTRPV1 receptors, indicating that the activation kinetics of RpTRPA5B are lower than dTRPA1-D and rTRPV1, as can be appreciated by channel ‘noise’ in the current traces (66), reflected also in the slower time course of activation of RpTRPA5B compared to the other thermosensors studied here. The maximum Po value reached experimentally in our study for rTRPV1 thermal activation is ∼1, in accordance with previous studies (49), however the maximum open probability of many ion channels is, in fact, typically lower than Po=1.0. Low threshold T-type voltage-activated channels (67), NMDA receptors (68), and thermoTRP channels like TRPM8 (69) all show experimental maximal open probabilities lower than Po=1.0. Likewise, voltage-gated Shaker K^+^ channel has an activity plateau at ∼ Po=0.7 (70). All these receptors have a great influence on the excitability displayed by the cells expressing them.

TRP channels are allosteric receptors, meaning that each stimulus (temperature, voltage, agonist) is detected by an independent module able to activate either partially or fully the channel. In the case of RpTRPA5B, temperature seems to be a partial activator given that the open probability is significantly lower compared to the other receptors. This observation is not a predictor that the channel may be open fully in response to different types of stimuli, or combination thereof (51). Additionally, RpTRPA5B presents slow activation kinetics, compared with the other receptors studied under the same conditions. There have been previous reports of other highly sensitive thermosensor receptors with slow activation kinetics such as hTRPV3. This channel exhibits a high temperature sensitivity, comparable to its homolog hTRPV1, but presents slower kinetics (51, 66). It is possible, like in the case of hTRPV3, that RpTRPA5B’s intrinsic molecular interactions influence the speed of the transitions between closed and open states (9), which would be interesting to test functionally, as it could potentially be explained by our observations of several structural differences in RpTRPA5B (Fig. 3).

From a thermodynamic point of view, many TRP ion channels are modulated by temperature and thus can integrate voltage and temperature allosterically (65). To disentangle these two properties, we specifically established the temperature sensitivity of the channel directly from the van’t Hoff plot, and not from the potential influence of temperature itself on the voltage activation process. Since we do not have evidence that the receptor can be activated by voltage, this allowed us to establish the thermodynamics of the temperature activation, completely independently from other stimuli sources, clearly supporting that RpTRPA5B is directly activated by temperature as the sole stimulus and belongs to a restricted category of thermoTRPs (8, 51). Finally, RpTRPA5B is also activated in a higher noxious range compared to known invertebrate thermoTRPs characterized thus far, including the fruit fly Painless and TRPA1 channels that mediate thermal nociceptive escape through larval mdIV neurons at temperatures above 40°C and 46°C, respectively (17), or Pyrexia channels that induce paralysis in adult flies upon exposure to 40°C (18). Altogether, RpTRPA5B is a temperature gated TRP receptor, with high-temperature sensitivity and activation responses to noxious heat stimuli *in vitro*. In mammals, only the vanilloid TRPV2 receptor contributes to such highly noxious (>52 °C) heat sensing (13, 52, 71). The expression platform implemented here has the potential to open up for comparative functional studies of TRPA5 orthologues as a potentially new class of highly noxious physiological sensors.

### Structural key features and new insights into TRPA5B’s pore and ankyrin domain

All *Rhodnius* and *Drosophila* ankyrin TRP monomeric structures were modelled with AlphaFold after validation with dTRPA1 (Fig. 3A). These models are snapshots of certain conformations and evidently do not reproduce the diversity of the different states a protein may adopt (72). Under default settings, AlphaFold provides a single state of otherwise highly dynamic proteins and is better at modelling backbone folding than individual sidechains. Taking these limitations into consideration, except for the orientation of the C-terminal helices, our results indicated that RpTRPA5B shares fundamental conserved regions and structural features with all modelled TRPA monomeric protein units (Fig. 3B), and support overall reliable secondary structure predictions close to tetrameric models and structures (Fig. 3A). We further ran pairwise comparisons of the monomeric structures of *RpTRPA5B* and Pyrexia, alternatively retained in hemipteran and dipteran genomes, respectively (Fig. 1). Interestingly, the two channels do not appear to occupy convergent homologous structural niches despite similarity in the ankyrin repeat domain (ARD, Fig. S7). Instead, RpTRPA5B appears to be generally closer in structure to Painless and Waterwitch in the transmembrane domain, while uniquely deviating in the specific details from modelled TRPA1 channels in the pore helices that flank the selectivity filter important for gating, and ARD previously suggested to contribute to thermosensitivity (73).

To start addressing the potential relevance of these differences in the pore and ARD, we first validated a truncated tetrameric model of dTRPA1, which aligned very confidently to the channel released structure in state 2 (PDB ID 7YKS) with a low RMSD for the twenty residues forming the upper and lower gate in the conductivity pore, even when compared to alignments of the same channel in state 1, especially if the Cα are considered (Fig. S8, Table S6). The modelling of the pore in RpTRPA5B is thus expected to be more reliable in the zones defined by the backbone, and interestingly appears narrower than the closed conformation of hTRPA1 (60), but with a wider lower gate (Fig. 3D-G. Fig S9). In contrast to the lower gate, which is wider due to the position of helix S6, the constriction at the selectivity filter is mainly determined by the orientation of a side chain (Glu914), which is less reliable and likely to be more dynamic. In hTRPA1, Gly914 in hTRPA1 is suggested to be important in gating, and lies in the location of the selectivity filter (60). It may seem peculiar that this highly conserved glycine residue is absent in RpTRPA5B, however a tetrameric model built simply inserting a Glycine residue in RpTRPA5B at that location causes significant deviations from the generic fold of TRPs including misplacement of helix S6 between adjacent monomeric units (Fig. S10). These observations suggest that the expected folding is likely properly maintained owing to compensatory co-evolving adjacent changes in the protein sequence.

Typically formed by repeats of 31 to 33 residue protein motifs that occur in tandem arrangement, N-terminal ankyrin repeat domains are known to be critical for a number of physiological processes such as ligand binding or protein-protein interactions and occur in a wide range of proteins including key sensory transducers of TRPA, TRPC, TRPN and TRPV channel families (74). Compared to fruit fly Pyrexia and other insect TRPA channels, RpTRPA5B not only displays a higher number of ARs but features longer loops, including between the third and the fourth ankyrin repeats, within the fifth ankyrin repeat, and between the fifth and the sixth ankyrin repeats, counting from the N-terminus. Another interesting feature is the disruption in the ankyrin repeat stacking between the fifth and the sixth ankyrin repeat in both *Rhodnius* and *Drosophila* Painless, which is not seen in *Drosophila* Pyrexia and RpTRPA5B. Although the potential impact of this conserved difference is currently unknown, we note that this breaking point coincides with the resolved N-terminal end of the recently reported structure of dTRPA1-A in state 2, which has been suggested to represent a temperature-sensitized, pre-opened conformation of the channel (58).

There is some evidence that the AR domain of some insect TRPAs may contribute to thermosensitivity, although more functional studies are needed prior to generalizing the role of ankyrin domains, and establishing correlations linking variation therein with global and specific mechanistic and thermosensitive properties. In particular, transfer of a part of the ARD from dTRPA1 (AR10-15) to hTRPA1, produced a heat-sensitive hTRPA1 (73), corroborating a contribution of this region to thermal activation sensitivity in the fruit fly dTRPA1(75, 76). In vertebrates, two regions of 6 ARs each in the snake thermosensitive TRPA1 (AR3-8; AR10-15) have been shown to revert the channel thermal sensitivity by conferring heat-sensitivity to a chimeric AR hTRPA1 (73). The temperature-dependent dynamics of the ARD have also recently been investigated in the TRPV1 channel, demonstrating that the ARD undergoes structural changes at similar temperatures that lead to TRPV1 activation, which suggested a potential role in the temperature-dependent structural changes leading to the channel opening (75). The N-terminus region of mosquito TRPA1 also seems to be quite critical for heat-sensitivity (19); however, there have been contradicting data for TRPA1, both from human and mosquito, arguing that additional regions controlling thermosensitivity are located outside the ARD (26, 77, 78). Altogether, with the understanding of limitations and constraints inherent to AlphaFold, these structural insights provide an interesting and relevant assessment of conserved key features of ankyrin TRPs for RpTRPA5B and underlie relevant structural novelties that may guide further functional studies in disentangling the proximate molecular determinants of the channel thermal and biophysical activation properties.

### Perspective on vector control strategies for *Rhodnius prolixus* considering plausible physiological roles for TRPA5

Sensory receptors in the same clades are often tuned to detect a stimulus over a discrete window of intensities, enabling the recognition of physiologically relevant cues over a wide dynamic range (1, 3, 79). The TRP Ankyrin family is an excellent example of this pattern as distinct, yet closely related channels, account for thermal responsiveness over a range from innocuous to noxious heat (4, 12). In addition, orthologous thermoTRPs often have different activation temperatures, and this has been postulated to reflect functional adaptive evolution to different optimal temperatures, coordinating thermoregulatory behaviours such as host-seeking, thermal avoidance, and tracking of optimal temperatures (80). Despite living in a wide range of ecological conditions, insects show overall little variability in maximum temperature that they can tolerate in an active state without inducing neural and physiological damage (40-50C) (81), except certain thermophilic ants that can forage above 50 °C for limited periods of time (82–84). The perception of environmental temperatures occurs through various organs and through the nervous system (83). RpTRPA5B is expressed broadly across tissues similarly to other insect thermoTRPs(18, 28), which together with functional validation of the channel *in vitro* activation by temperature, is in line with a possible physiological role in thermosensation. To investigate its physiological role in heat detection, it will be interesting to map the cellular location of RpTRPA5B in peripheral sensory neurons including neural populations in the central nervous system, as shown in dTRPA1-D or *Pyrexia* channels (18, 22), channels associated with physiological thermotolerance in the fruit fly (83).

Variable environmental temperatures are extremely common in natural environments of small insects with low thermal capacity, thus, detecting and avoiding heat is critical to prevent injury. Temperature distributions vary widely for natural objects. For instance, dry and moderately gray-coloured or dark objects such as tree bark or rocks easily reach temperatures above 50°C (84). If the humidity level is high, and radiative cooling of the sky is not effective, the same objects can reach temperatures above 60°C in the full sun. For example, temperatures of dry leaf substrates on the ground can exceed 50°C in full sun since they do not undergo evaporative cooling, which would typically prevent a leaf’s surface temperature from going above 40°C. In lab-simulated natural environments and in field thermal imaging studies, insects can reach 60°C under full sun with high humidity in as little as 15 seconds (84). Considering that *Rhodnius* adults are about 3 cm in length and dark coloured, with a small thermal capacity, and that they typically inhabit tropical environments with high humidity, they are likely to rapidly reach temperatures above 60°C if exposed to full sun, further suggesting that TRPA5 may mediate noxious heat avoidance in this species. Although the physiological and behavioural role of RpTRPA5B will need to be examined in detail, the discovery of the activation range of TRPA5B opens avenues for exploring the convergent evolution of noxious heat sensing by TRPA5 and for assessing whether it is also relevant for both inner temperature regulation and warm substrate avoidance.

ThermoTRPs are polymodal sensors of physical and chemical stimuli (79). For example, channels in the insect TRPA1 and HsTRPA clades are typically activated by allyl isothiocyanate (AITC) and various plant-derived chemicals such as carvacrol and citronellal (23, 28, 29). However, characterized receptors of noxious heat in insects such as Pyrexia and Painless do not exhibit chemical sensitivity to electrophiles (17). From an evolutionary perspective, a role for the TRPA5 clade as polymodal sensor exhibiting heat and chemical sensitivity is plausible. Live *Rhodnius* individuals treated with capsaicin, the vanilloid pungent extract of chili peppers, were recently shown to have impaired orientation towards a thermal source (85). Notably, this compound can directly activate the mammalian TRPV1 receptor independent of temperature, and the mammalian noxious temperature receptor, TRPV2, when bearing only four mutations (3, 15). Other than capsaicin, both TRPV1 and TRPV2 are readily activated by additional vanilloid compounds such as resiniferatoxin, an active compound from the cactus *Euphorbia resinifera* used for medicinal purposes and other plant-derived compounds that act as chemical agonists (86). Findings of botanical compounds triggering chemical activation of RpTRPA5B combined with *in vivo* behavioral exposure studies, may thus contribute to uncovering new classes of natural repellents potentially co-mediating heat-avoidance. This would be significant, not only in *Rhodnius* but also for other triatomines and hemipteran vectors sharing a close TRPA5B orthologue such as the bed bug, *Cimex lectularius*.

The findings of an insect ankyrin TRPA5 ion channel sensitive to noxious temperature *in vitro*, highlights independent evolutionary origins of the molecular transduction of thermal stimuli in insects, while simultaneously opening the door for further pharmacological studies of TRPA receptors in triatomine vectors.

## Material and Methods

### Phylogenetic analyses

Amino acid sequences of insect TRPA channels from the Anoplura (sometimes included under Psocodea or Phthiraptera), Coleoptera, Diptera, Hemiptera, Hymenoptera, Isoptera and Lepidoptera insect orders were retrieved from the InsectBase repository (87), FlyBase version FB2020_03 (88), VectorBase (https://www.vectorbase.org), BeeBase (89), NCBI-blast (90), EnsemblMetazoa (https://metazoa.ensembl.org), the i5k Workspace@ NAL (91) and OrthoFinder (92). The TRP sequences from insect model systems including *Drosophila melanogaster, Tribolium castaneum, Bombyx mori, Apis mellifera* and *Rhodnius prolixus* were used as templates to mine and curate orthologous TRP ORF sequences from annotated insect genomes and transcriptomes. To classify the uncharacterized TRPs, amino acid sequences were aligned using MAFFT (93), and Maximum-Likelihood phylogenetic trees were inferred in IQ-TREE v1.6.11 using ModelFinder (Ultrafast Bootstrap, 1000 replicates), using a best-fit model JTT+F+I+G4 measured by the Bayesian information criterion (BIC) (94–96). The phylogenetic trees were visualized, rooted at mid-point and annotated in R V3.6.3 using the ggtree package (97) and Evolview (98). The accession numbers are listed in Table S1.

### TRPA5 gene annotation and tissue expression

We collected Illumina read data from *R. prolixus* tissue libraries published in the Sequence Read Archive (SRA) at NCBI under Bioproject accession numbers PRJNA281760/SRA:SRP057515 (antennal library from larvae, female adult and male adult) (43), PRJEB13049/SRA:ERP014587 (head library), and PRJNA191820/SRA:SRP006783 (ovary and testes library). We performed low-quality base trimming and adaptor removal using cutadapt version 1.16 (99) and aligned the trimmed read pairs against the *R. prolixus* assembly version RproC3.0.3 (retrieved from VectorBase.org) genome using HISAT2 version 2.2.0 (100). The existing annotation was used to create a list of known splice sites using a python script distributed with HISAT2. We used StringTie version 2.1.3b (101) with the *conservative* transcript assembly setting to improve the annotation, reconstruct a non-redundant set of transcripts observed in any of the RNA-Seq samples, and compute expression estimates.

We applied Trinotate version 3.2.1 (102) to generate a functional annotation of the transcriptome data. In particular, the functional annotation of TRP genes for which the initial genome annotation was absent or incomplete (i.e *TRPA5, Nan, Pain*) were localized in Trinotate annotation followed by validation using the Apollo gene browser (103). All TRP gene identifiers are presented in Table S2.

The alignment BAM files were used to estimate transcript abundance using StringTie together with our improved annotation. The abundance tables from StringTie were imported into R using the *tximport* package (104), which was used to compute gene-level abundance estimates reported as FPKM. We used the R package *pheatmap* to visualize the expression level of TRP genes.

### Monitoring of TRPA5B expression levels by quantitative PCR

Live adults of *R. prolixus* were obtained from BEI Resources (USA). Female antennae, rostrum, legs, heads (minus antenna and rostrum), and bodies (thorax minus legs + abdomen) were dissected and pooled from 15 individuals in DNA/RNA shield reagent (Zymo) and stored at -20°C until further processing. Total RNA was isolated using the Monarch RNA extraction procedure (New England Biolabs), including tissue grinding in liquid nitrogen and a DNAse I step. cDNAs were synthesized using the GoScript cDNA synthesis procedure (Promega) prior to concentration assessment using the Qubit High sensitivity DNA kit (Invitrogen). Two gene-specific primer (GSP) sets were designed for *Rhodnius* Actin (Genbank acc. Nr. EU233794.1) and TRPA5B using Primer3 version 2.3.7 in Geneious (105) (Table S5). Each primer set was initially validated by calculating standard curves from serial dilutions of template cDNA (2 ng/μL to 0.25 ng/μL) and primer mix (5 to 0.25 μM) with choosing amplification efficiencies (E) between 95 and 100%. qPCR amplification products from initial runs were additionally checked on 2% agarose gels to verify the correct amplicon sizes and the absence of primer dimers. As a final validation, qPCR products were purified using Exo-SAP (Fermentas) prior to Sanger sequencing to ensure product amplification specificity. Quantitative PCR reactions were then run in three technical replicates on a CFX384 Real-Time PCR system (Bio-Rad) with quantification and dissociation curves analyses performed for three independent experiments using the CFX Maestro Software 2.3 (Bio-Rad). Each five-microliter reaction contained 2.5 μL 2x SsoAdvanced Universal SYBR Green Supermix (Biorad), 0.25 ng cDNA and 0.125 μM primers. Cycling conditions were as follows: 95°C for 2 min, 39 cycles of 95°C for 10 s, 60°C for 10 s followed by a dissociation curve analysis from 65.5°C to 89.5°C with gradual heating at 0.6°C/s. Relative log-fold expression levels were normalized per tissue type against the reference gene and calibrated relative to Antennae (log fold expression = 1)

### Alpha-fold modeling and DALI analyses

Monomer structures of *Rhodnius* TRPA1, *Rhodnius* Painless, *Rhodnius* Waterwitch, *Rhodnius* TRPA5B, *Drosophila* TRPA1-D, *Drosophila* Painless, *Drosophila* Waterwitch and *Drosophila* Pyrexia were generated using AlphaFold2 with amber relaxation activated (55) on Colab’s server (57). The sequence of dTRPA1 isoform D used for the AlphaFold prediction is identical to the sequence of the determined structure (isoform A) except for the first 97 amino acid residues, in which the five last residues correspond to the first five in the resolved N-terminal region in 7YKR. Using the same tool, a tetrameric model of residues 477-1153 of *Drosophila* TRPA1-D corresponding to PDB ID 7YKS was made. No template was used in these predictions. To model the Rhodnius TRPA5B tetramer, due to limitations in computational power, the transmembrane region (residues 608-1078) was modelled first, and then used as a custom template to model a monomer of residues 42-1078. The first 41 residues and the C-terminal of the monomers from residue 1079 were disordered and truncated to avoid clashes when assembling the tetramer. A tetramer was assembled of four copies of the monomer by aligning them to each of the chains of the truncated transmembrane tetramer in PyMOL (106). The monomer models were compared with pairwise structural alignment using the Dali server (107). The PDB files are provided as source datafiles.

### Molecular cloning

Antennae from twenty Rhodnius adult individuals were obtained from a laboratory culture (Orchard lab, University of Toronto Mississauga, Canada) and stored in DNA/RNA Shield^TM^ reagent (Zymo Research). Tissues were disrupted in Trizol using a Premium Multi-Gen 7XL Homogenizer (PRO Scientific) and RNA was subsequently extracted using the Direct-zol RNA kit (Zymo Research), including a DNAse step to remove genomic DNA contamination. cDNA was synthesized from 1ug Total RNA using the GoScript^TM^ Reverse Transcriptase kit (Promega) and random hexamers following the recommended manufacturers’ protocol. RNA and cDNA qualities were verified using a Nanodrop (Nanodrop 2000/2000c UV-vis spectrophotometer, Thermo Scientific) and quantified using a Qubit Fluorometer (ThermoFisher). The coding regions of Rhodnius *Rp-TRPA5B* was amplified from antennal cDNA using gene-specific primers designed based on Rhodnius full length TRP sequences (36) and containing unique restriction sites (Table S6). PCR reactions were performed in a Veriti™ Thermal Cycler (ThermoFisher) using the Advantage® 2 PCR Kit (Takara Bio) in a touchdown cycling program as follows: 95°C for 2 min, 16 cycles of 95°C for 30 sec, 68°C for 1 min (-0.5°C/cycle), 68°C for 4 min followed by 20 cycles of 95°C for 30 sec, 60°C for 1 min, 68°C for 4 min, and a final step at 68°C for 10 min. Amplification products were analysed by electrophoresis, and fragments of expected size were excised from the gel, purified using the Monarch® DNA gel extraction kit (NEB) and subjected to Sanger Sequencing for ORF sequence-verification prior to codon-optimization at Genscript and subcloning. For the rat rTRPV1 and the fruit fly dTRPA1-D, gene specific primers (Table S6) were used to amplify the ORF including suitable flanking restriction sites prior to gel purification and double restriction digestion. The digested PCR products were gel purified and ligated in an expression cassette containing the human cytomegalovirus (CMV) immediate early promoter and engineered to include a C-terminal tag by the monoclonal antibody FLAG epitope sequence (DYKDDDDK), followed by a Ser-Gly-Ser linker peptide, a T2A peptide sequence (EGRGSLLTCGDVEENPG) and the coding region of the cytoplasmic fluorescent marker protein mRuby2 (48, 50). The ligation mixtures were used to transform *Stbl3* competent *E. coli* cells (ThermoFisher) using standard protocols. Plasmid DNAs were purified using the Qiaprep spin Miniprep (Qiagen) and verified by Sanger sequencing using internal gene-specific and vector primers to ensure overlapping sequence information in both forward and reverse directions. High yield pure plasmid DNA preparations were subsequently obtained from 100 mL overnight LB broth cultures using the endo-free ZymoPURE™ II Plasmid Midiprep Kit (Zymo Research, USA).

### Transient HEK293T cell expression

Plasmid DNAs clones from TRP cDNAs were transiently expressed in HEK293T cells to optimize expression conditions via mRuby2 visualization and western blot analysis prior to whole cell patch clamp recordings. HEK293T cells were seeded at a density of 0.6 x 10^6^ cells on day 0 in 60 mm culture dishes (ref 25382-100, VWR) in DMEM High Glucose, GlutaMAX (Life Technologies) supplemented with 10% FBS (Seradigm Premium, VWR, USA). For each transfection, lipid complexes containing 2.5 μg DNA: 10 μL L2000 (Life Technologies) mixed in Opti-MEM I Reduced Serum (Life Technologies) were added dropwise to the cells at 50% confluency (1.2 x 10^6^ cells). The culture medium was exchanged with new DMEM/FBS medium six-hours post-transfection. Cells were incubated at 37°C in a humidified HERAcell 150i incubator (Thermo Scientific) with 5% CO_2_.

### Biochemistry

For whole-cell TRP expression analysis, cells were harvested 72h post-transfection; the medium was decanted, cells were collected in 2mL cold D-PBS, centrifuged for 5 min at 4,000 rpm at 4°C and then the supernatant was discarded. The cell pellet was gently suspended in 50 μL cold Ripa lysis buffer (Thermo Scientific) supplemented with 1% Triton-X100 (Sigma-Aldrich) and complete EDTA-free protein inhibitors (Sigma-Aldrich). Cell membranes were lysed for 1h at 4°C with gentle rotation on a sample homogenizer, and cell debris were collected by centrifugation at 4°C for 15 min at 13,000 rpm. The crude protein lysate concentration was quantified by bovine serum albumin (BSA) (Sigma-Aldrich) and 25 μg crude extract was loaded on NuPAGE™ 3-8% Tris-Acetate gels (ThermoFisher) and transferred to a polyvinylidene difluoride membrane on a TurboBlotTransfer system (Bio-Rad Laboratories). The membranes were blocked with 5% milk (Bio-Rad) in Tris-buffered saline containing 0.1% Tween 20 (TBST, Bio-Rad) and incubated overnight with aFLAG antibody 1:2,500 (GE Healthcare) on a gently rocking platform at 4°C. After washing with TBST the membranes were incubated for 1h at ambient temperature in the dark with horseradish peroxidase (HRP) ECL anti-mouse conjugated antibody (Amersham, USA) diluted in 5% milk in TBS-Tween at 1:2,500. Membranes were rinsed in TBST and revealed using the SuperSignal West Femto (Thermo Scientific) and imaged on a ChemiDoc system (Bio-Rad Laboratories).

For membrane surface expression, the plasma membrane expression of RpTRPA5B channels was assessed using the Pierce Cell surface Protein isolation kit (Thermo Scientific). On day 0, four T75 cm^2^ flasks were seeded with 1 x 10^6^ HEK293T cells and incubated at 37 °C. Forty hours later, each flask was transfected with lipid complexes containing 48 μg endo-free plasmid DNA and 96 μl Lipofectamine 2000 diluted in Opti-MEM serum and incubated at 30°C. 72 hours post-transfection, cells were gently washed with ice-cold PBS, labeled with Sulfo-NHS-SS-Biotin, and harvested following the manufacturer’s protocol. Cells were lysed on ice for 30 min in the manufacturer’s lysis buffer supplemented with 0.5% Triton-X100 and complete EDTA-free protein inhibitors (Sigma-Aldrich), with gentle 5s vortexing every 5 min, and two 5x-1s sonicating bursts on ice. Following centrifugation, the cell lysate was bound to NeutrAvidin agarose resin and gently mixed for 60 min at ambient temperature on a platform rotator. The membrane-bound fraction was eluted with 50mM Dithiothreitol in SDS-Sample buffer (62.5 mM Tris/HCl pH6.8, 1% SDS, 10% Glycerol) and then placed on ice. For Western Blot analysis, 50 µg of the membrane protein eluate fraction quantified by BSA were diluted to 32 μl in lysis buffer containing loading Laemmli buffer (Bio-Rad) supplemented with 10% 2-mercaptoethanol. Sixteen μl (25 µg) of the homogenized protein-loading buffer sample were loaded in duplicates on a NuPAGE™ 3-8% Tris-Acetate gel (ThermoFisher) to be probed separately with FLAG and ATPase antibodies. Proteins were separated by electrophoresis for 3h at 80V at 4°C, then transferred to a polyvinylidene difluoride membrane on a TurboBlotTransfer system (Bio-Rad Laboratories). The membranes were blocked in parallel with 5% milk (Bio-Rad) in Tris-buffered saline containing 0.1% Tween 20 (TBS-T, Bio-Rad) and incubated overnight on a gently rocking platform at 4°C with aFLAG antibody 1:2,500 (ab213519, Abcam) or with Anti-Sodium Potassium ATPase antibody 1:2,500 (ab76020, Abcam) diluted in 5% milk. After three washes with TBST, the membranes were incubated for 1h at ambient temperature in the dark with HRP ECL anti-mouse conjugated antibody (Amersham, USA) at a 1:2,500 dilution in 5% milk/TBST. Membranes were rinsed in TBST and revealed using the SuperSignal West Femto (Thermo Scientific) and imaged on a ChemiDoc system (Bio-Rad Laboratories).

### Temperature control using a laser system

We used a manual patch-clamp station (Axopatch 200, Molecular Devices) equipped with a fiber-delivered laser system to record temperature-activated currents under a precise voltage-clamp control. The setup was modified after Yao et al (2009) (108) and takes advantage of water’s IR absorption band to generate rapid temperature jumps from RT to high temperatures. It combines an infrared diode laser (λc =1460 nm (+/-20 nm), Output power = 4.8 watts) (Seminex Inc.) coupled with a 100-um optical fiber with a striped tip (ThorLabs, Inc.) as the controllable heat source. Two independent micromanipulators allowed us to precisely align the relative positions of the patch-clamp electrode and the fiber on a single cell (Fig. 3, Fig. S9). To calibrate the optic fiber position with respect to the patch pipette we used a visible laser (Fig. S9). Marks on the computer screen were used to keep the position of the fiber and the pipette consistent for the different experiments. Cells under whole-cell voltage-clamp control were held at -30mV during the experiment. To program fast pseudo-transient temperature changes, the patch pipette current was used to read the temperature changes in real-time as the feedback to the laser diode controller (LDC-37620, ILX Lightwave) to perform proportional-integral-derivative (PID) control of the driving current of the laser diode (*see Supplemental Methods*). This laser-heating setup provides a rapid and precise heating rate on the order of 50°C within tens of milliseconds, essential to provide both adequate temporal resolution and controllable steady-state temperatures in the range of 35°C to 70°C to analyze the channel activation. Fig. S9A shows constant temperature steps were achieved with a rising time constant of 34.2±3.3 ms, independent of the laser power. The temperature jump associated with successive current pulses is precisely calculated by running an open pipette calibration following the same current sequence at the end of each run (Fig. 3D).

### Temperature calibration

We used the resistance of the open pipette to measure the temperature jump magnitudes following the equation T={1/T_0_ – R/E_a_ x ln(I/I_0_)} - 1, where R is the gas constant, T_0_ and I_0_ are respectively room temperature and the corresponding electrode current at room temperature. The activation energy (E_a_) of the system corresponds to 3.84 kcal/mol as was established by Yao et al. (2010) for the pair of solutions used in the recordings (108). The equation describes the change in ion motility as a function of temperature changes in the system. The current change was used as a feedback signal for a laser-diode controller software coded in Labview that uses a proportional-integral-derivative (PID) control algorithm. To account for the variability in the diameter between the different patch pipettes used in different experiments, the instrument was calibrated before each experiment to assure comparable temperature jumps in each experiment, adjusting the diode power outputs to the desired temperature accordingly.

### Whole cell patch-clamp recordings

Cells were seeded at low density in a 30 mm culture dish (VWR) containing round glass coverslips 48h post-transfection (Table S7; Fig. 3). Cells were first rinsed with D-PBS at room temperature, trypsinized with 0.5 mL Accutase (Stemcell Technologies) and suspended in 4.5 mL pre-warmed DMEM-FBS medium. Two hundred microliters of this cell suspension were mixed with 1.8 mL pre-warmed DMEM-FBS medium, dispensed drop wise in the culture dish, and incubated for 24h at 30°C. In a typical experiment, one glass cover slip was gently retrieved from the culture dish using sterile forceps, rinsed with a recording solution using a Pasteur pipette and placed in the recording chamber. The fluorescence of mRuby-expressing cells was monitored to select bright, healthy, isolated cells for whole-cell patch clamping. Experiments under the whole-cell configuration were carried out 72h after induction. The electrodes were fabricated with borosilicate capillaries, using a horizontal micropipette puller P-1000 (Sutter Instrument, Novato, CA, USA), and polished using a microforge (Narishige, Japan) to a final diameter between 2-4 um. The internal electrode was filled with the following solution in mM: 126 CsCl, 10 EGTA, 10 HEPES, 1 EDTA, and 4 MgATP. The extracellular recording solution contained 2 mM CaCl_2_, 10 mM HEPES, 140 mM NaCl, pH 7.2 (adjusted with NaOH). The electrode resistance ranged between 2-4 MΩ, and the Vjp was estimated at ∼18 mV for the recordings. The current traces were amplified using a MultiClamp 700B amplifier (Molecular Devices, Sunnyvale, CA, USA). The amplified electrical signals were acquired with a Digidata 1440A using the pClamp10 software (Molecular Devices, Sunnyvale, CA, USA). Series Resistance (Rs) was compensated in 80%, as well as the fast and slow capacitive components of the current. The current density was fitted to the following Boltzmann function:

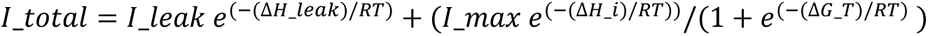

Whereby the first term Δ*H_leak* is the enthalpy change of the leak current. The second term accounts for the channel activity, with Δ*G* = Δ*H* − *T*Δ*S* is the free energy change involved in the closed-open reaction, and Δ*H*_*i* accounts for the linear temperature dependence of the ionic conductivity and leakage current (108). The corrected temperature current density (*I*) was used to calculate the equilibrium constant from the relative fraction of the channel in the open conformation (Po), assuming a two-state model, where Po = I/Imax.

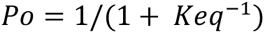

where ln(K_eq_) = -(ΔH/RT) + (ΔS/R). Thus from the van’t Hoff plots ln(Keq) vs 1/T, the enthalpy and entropy associated with the channel opening can be obtained (51). The channel temperature activation parameters are obtained from the normalized current (Gmax or I/Imax) following methods for both cold-and heat-activated ion channels (49, 69). Normalization of the maximum amplitude against an agonist does not influence the thermodynamic parameters as determined from the normalized current.

## Acknowledgments

The authors acknowledge Andrew Allen for helpful discussions on TRP biochemistry, Xi-Shi for early advice with HEK cell culture, Rhiannon Macrae for her valuable guidance, Feng Zhang for his support and for giving access to resources at the Broad, Ian Orchard for providing *Rhodnius prolixus* antennae, BEI resources (http://www.beiresources.org/) and Ellen M. Dotson for providing Rhodnius adults, Rachel Gaudet for providing a rTRPV1 plasmid template and Pengyu Gu for providing a *Drosophila* dTRPA1-D (A10a, flybase PG) plasmid template. This work was supported by a Mind Brain Behavior Interfaculty grant to NEP and MAL, an Alice and Knut Wallenberg fellowship at the Broad Institute of MIT and Harvard to MAL, National Science Foundation PHY-1411123 and PHY-1411445 to NEP and NY respectively, funding from Formas 2017-01463 to UJ, from the Carl Tryggers Foundation (CTS 21:1309) and the Swedish Research Council (VR2020-05107) to MAL. JML is supported by a MISU grant from the FRS-FNRS (F.6002.22).

## Data availability statement

A supplementary information file containing supplementary methods and supplementary figures is provided as an accompanying PDF file. Supplementary tables are grouped in a supplemental excel document. A source data document accompanies the submission that lists all source data usage including Figshare links.

## Code availability statement

Custom codes used to generate phylogenies and graphs are available in the source data file (Figshare links).

## Contributions

MAL and NEP conceived the study. MAL administered the project, coordinated all experiments and data analysis. MAL, DBN, CCT, NFY designed the study with input from JQP and NEP. CCT and NFY implemented the laser setup with input from DBN. MAL, DBN, CCT and WAVM performed experiments. MAL conducted and analyzed the molecular aspects of the project. DBN conducted and analyzed the *in vitro* electrophysiology. JML conducted the genomic and transcriptomic analyses. BW and UJ conducted the structural and modelling analyses. MAL, DBN, UJ, BW and JML provided visualizations. NEP, NFY and JQP provided resources. MAL and DBN wrote the manuscript draft, and all authors contributed to the writing and editing of the manuscript.

