## Supplemental Information for "Convergent evolution of noxious heat sensing by TRPA5, a novel class of heat sensor in *Rhodnius prolixus*"

**This PDF file includes:**

1. **Supplementary Methods**


1.1. Supplementary Method 1: PID control

1.2. Supplementary Method 2: Open current pipette measurements

1.3. Supplementary Method 3: Fiber preparation


2. **Supplementary Figures**


2.1. Supplementary Figure 1: Phylogeny of insect TRPA channels

2.2. Supplementary Figure 2: Phylogeny of insect TRPA5 channels

2.3. Supplementary Figure 3: qPCR analysis of RpTRPA5B

2.4. Supplementary Figure 4: TRPA5B whole cell and surface expression

2.5. Supplementary Figure 5: Validation with rTRPV1 and dTRPA1-D

2.6. Supplementary Figure 6: Open pipette temperature calibration and optic fiber positioning

2.7. Supplementary Figure 7: DALI analysis

2.8. Supplementary Figure 8: Comparison of experimentally determined structure of dTRPA1 in state-2 (PDB ID 7YKS) and dTRPA1 tetrameric AlphaFold model

2.9. Supplementary Figure 9: Distances between corresponding residues in the upper and lower gates of dTRPA1, hTRPA1 and RpTRPA5B.

2.10. Supplementary Figure 10: Comparison of 6V9Y and RpTRPA5


**Other Supplementary Materials for this manuscript include the following:**

**1. Grouped in a single separate .xls file:**

- Supplementary Table 1: TRPA gene accession numbers

- Supplementary Table 2: *Rhodnius prolixus* TRP gene identifiers
- Supplementary Table 3: qPCR data for RpTRPA5B tissue expression analysis
- Supplementary Table 4: Cell density for whole patch recordings

  Supplementary Table 5: Ankyrin repeats in monomeric structure models of TRPAs
- Supplementary Table 6: Comparison of residues in the upper and lower gate between the AlphaFold tetrameric model and determined structures of dTRPA1
- Supplementary Table 7: Oligonucleotide primer sequences

**2. In a separate word document, a list of all source data for each figure, with corresponding figshare links.**

#### Supplementary Methods

##### 1.1. Supplementary Method 1: PID control

A PID (proportional-integral-derivative) control is the typical way to adjust the output according to the input reading in real-time without knowing most of the environmental parameters. The idea is described in the following function: $Output(t) = K_p Err(t) + K_i \int_0^t Err(\tau) d\tau + K_d \frac{dErr(t)}{dt}$

$$Err(t) = Input(t) - Setpoint$$

with $Output(t)$ being the laser power, $Input(t)$ being the temperature, and $Setpoint$ being the desired value. The $Output(t)$ is determined by $Err(t)$, which is the difference between $Input(t)$ and $Setpoint$.

There are 3 terms in this equation, the first term is the proportional term. This term varies linearly with $Err(t)$. For instance, when the temperature reaches the setpoint, this term decreases, and when the temperature exceeds the setpoint, this term becomes negative to bring the temperature back to the setpoint. The second term is the integral term. This term provides a gradually increasing offset and this offset will stabilize when the temperature stabilizes to setpoint, where $Err(t) \to 0$. The third term is the derivative term. This term estimates the required change of output by watching the inertia of $Err(t)$. For instance, when the temperature reaches the setpoint, the proportional term gives 0 and the integral term gives a stabilized value, but if the temperature is still increasing, this term will decrease the output to prevent the temperature from exceeding the setpoint in the next time interval.

##### 1.2. Supplementary Method 2: Open pipette current measurements

The temperature is measured by monitoring the current through an open patch pipette, which means there are no cells but only water, assuming the thermal property of the cell is the same as water. First, a series of open-pipette measurements with different laser powers and power waveforms is performed and used to calculate the temperature evolution from real-time current in the patch-clamp recording pipette. The patch-clamp experiment is conducted by applying the same laser powers and waveforms to the cell.

##### 1.3. Supplementary Method 3: Fiber preparation

The attenuation coefficient of water at the wavelength of 1460 nm is ~3000 $cm$^(−1) which means the absorption of the laser power through a 100$nm$ thick water layer is over 40%. Using the fiber above the water surface, the laser would thus deliver most of its power in the upper water layer. The temperature of water above the targeted cell would be much higher than the cell temperature itself, and the temperature change would not be confined to a single cell due to conduction and convection in the water. To resolve this, we cut the fiber with a diamond blade under a stereomicroscope and stripped the fiber which allowed us to place the fiber tip in the water layer directly above the cell in the recording chamber (Fig. S6). The relative position of the fiber tip and cell was adjusted using patch-clamp rig microscope.

### 2. Supplementary Figures

#### 2.1. Supplementary Figure 1: Phylogeny of insect TRPA channels

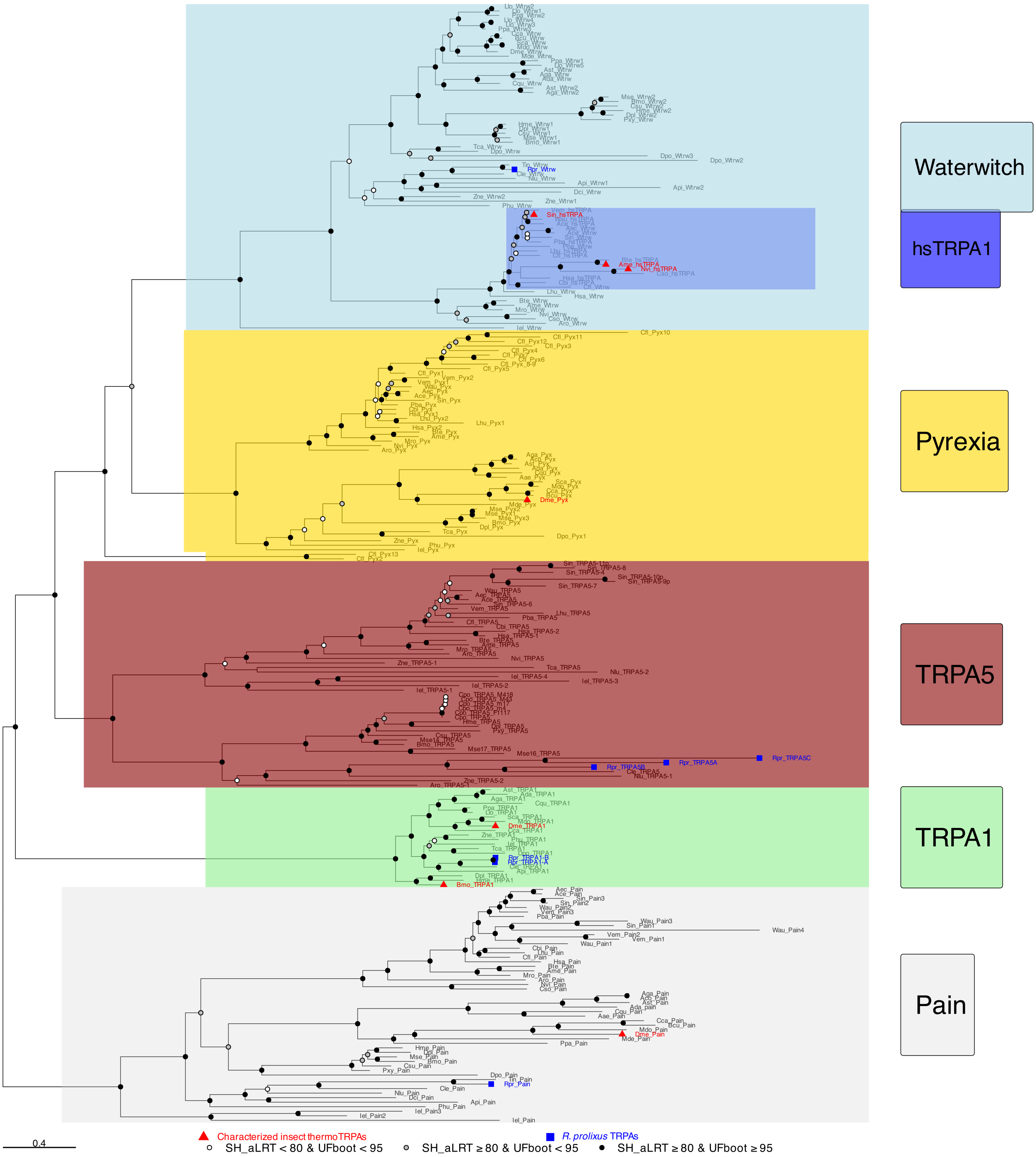

**Figure S1. Phylogeny of insect TRPA channels.** The Maximum-likelihood phylogeny of amino acid sequences includes representative insect ion channel members of the TRPA1, Pain, Pyrexia, hymenoptera-specific TRPA (hsTRPA), TRPA5 and Waterwitch (Wtw) subfamilies. Accession numbers are listed in Supplementary Dataset, Table S1. The tree was inferred in IQ-TREE v1.6.11 using ModelFinder, Ultrafast Bootstrap (UFboot), 1000 replicates, using a best-fit model JTT+F+I+G4 measured by the Bayesian information criterion (BIC). Branches were assigned Shimodaira-Hasegawa approximate likelihood ratio test (SH-aLRT) and UFboot supports. The tree was visualized in Rstudio (2021.09.2) using ggtree (Source data Figure S1). Species represented are **Anoplura**: *Pediculus humanus* (Phu, head louse), **Coleoptera**: *Tribolium castaneum* (Tca, red flour beetle), *Dendroctonus ponderosae* (Dpo, Mountain pine beetle), *Photynus pyralis* (Ppy, Common Eastern Firefly), *Agrilus planipennis* (Apl, Emerald ash borer), *Anoplophora glabripennis* (All, Asian longhorned beetle), *Aethina tumida* (Aut, Small hive beetle), *Onthophagus taurus* (Ota, bull dung beetle), **Diptera**: *Drosophila melanogaster* (Dme, fruitfly), *Mayetiola destructor* (Mde, Hessian fly), *Culex quinquefasciatus* (Cqu, Southern house mosquito), *Anopheles stephensi* (Ast), *Anopheles gambiae* (Aga), *Anopheles darlingi* (Ada), *Bactrocera cucurbitae* (Bcu, Melon fly), *Ceratitis capitata* (Cca, Mediterranean fruit fly), *Musca domestica* (Mdo, housefly), *Lutzomyia longipalpis (Llo, Sandfly), Phlebotomus papatasi* (Ppa, Sandfly), *Stomoxys calcitrans* (Sca, Barn fly), **Hemiptera:** *Acyrthosiphon pisum* (Api, pea aphid)*, Nilaparvata lugens* (Nlu, Brown planthopper), *Diaphorina citri* (Dci, Asiatic citrus psyllid), *Cimex lectularius* (Cle, Bed bug)*, Triatoma infestans* (Tin, Winchuka) *Rhodnius prolixus* (Rpr, Kissing bug)*,* **Hymenoptera:** *Apis mellifera* (Ame, Western honeybee) *, Bombus terrestris* (Bte, Buff-tailed bumblebee)*, Megachile rotundata* (Mro, Alfalfa leaf cutting bee), *Nasonia vitripenis* (Nvi, Jewel wasp), *Harpegnathos saltator* (Hsa, Jumping ant)*, Linepithema humile* (Lhu, Argentine ant), *Campanatus floridanus* (Cfl, Florida carpenter ant), *Pogonomyrmex barbatus* (Pba, red harvester ant), *Atta cephalotes* (Ace, Leafcutter ant), *Acromyrmex echinatior* (Aec, Panamian leafctutter ant), *Solenopsis invicta* (Sin, Red imported fire ant)*, Vollenhovia emeryi* (Vem, ant), *Athalia rosae* (Aro, Turnip sawfly), *Cerapachys biroi* (Cbi, Clonal raider ant), *Ceratosolen solmsi* (Cso, Fig wasp), *Wasmannia auropunctata* (Wau, Electric ant), **Isoptera:** *Zootermopsis nevadensis* (Zne, Dampwood termite), **Lepidoptera**: *Bombyx mori* (Bmo, Silkmoth)*, Chilo suppressalis* (Csu, Asiatic rice borer), *Danaus plexippus* (Dpl, Monarch butterfly), *Heliconius melpomene* (Hme, Postman butterfly), *Plutella xylostella* (Diamondback moth), *Manduca sexta* (Mse, Tobacco hornworm) *Cydia pomonella* (Cpo, Codling moth), **Odonata**: *Ischnura elegans* (Iel, Bluetail damselfly).

#### 2.2. Supplementary Figure 2: Phylogeny of insect TRPA5 channels

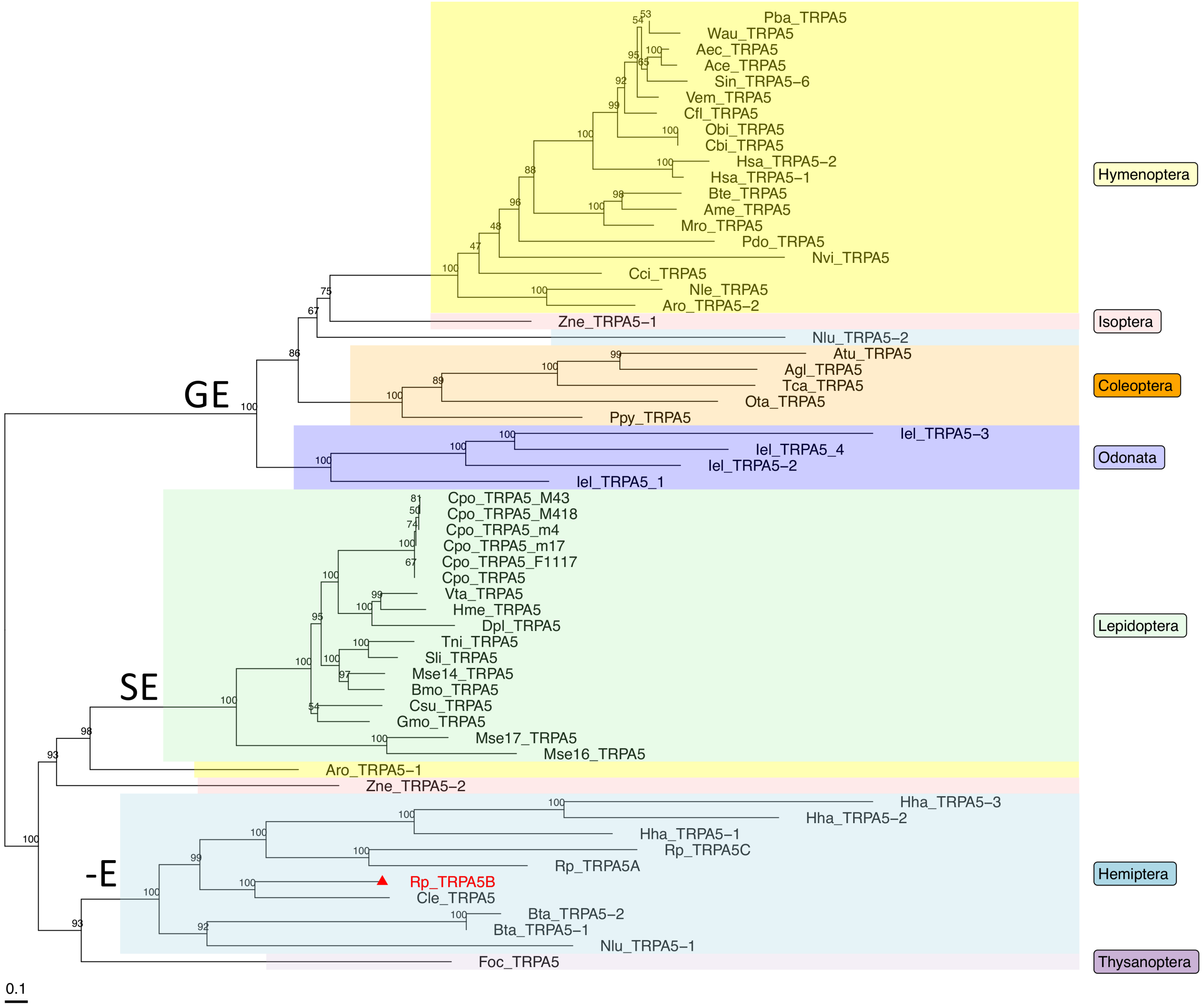

**Figure S2. Phylogeny of insect TRPA5 channels.** TRPA5 channels are present across insect Orders but Diptera. The Maximum-likelihood phylogeny of amino acid sequences includes representative ion channel members of the TRPA5 subfamily. The tree was inferred in IQ-TREE v1.6.11 using ModelFinder, Ultrafast Bootstrap (UFboot), 1000 replicates, using a best-fit model JTT+F+I+G4 measured by the Bayesian information criterion (BIC). Branches were assigned Shimodaira-Hasegawa approximate likelihood ratio test (SH-aLRT) and UFboot supports. The tree was visualized in Rstudio (2021.09.2) using ggtree (see Source data Figure S2). Species abbreviations are as follows: **Coleoptera:** *Tribolium castaneum* (Tca, red flour beetle), *Photynus pyralis* (Ppy, Common Eastern Firefly), *Anoplophora glabripennis* (Agl, Asian longhorned beetle), *Aethina tumida* (Aut, Small hive beetle), *Onthophagus taurus* (Ota, bull dung beetle); **Hemiptera**: *Bemisia tabaci* (Bta, Silverleaf Whitefly), *Cimex lectularius* (Cle, Bed bug)*, Halyomorpha halys* (Hha, Brown marmorated stinkbug), *Nilaparvata lugens* (Nlu, Brown planthopper), *Rhodnius prolixus* (Rp, Kissing bug); **Hymenoptera**: *Apis mellifera* (Ame, Western honeybee) *, Athalia rosae* (Aro, Turnip sawfly), *Bombus terrestris* (Bte, Buff-tailed bumblebee)*, Megachile rotundata* (Mro, Alfalfa leaf cutting bee), *Nasonia vitripennis* (Nvi, Jewel wasp), *Harpegnathos saltator* (Hsa, Jumping ant)*, Linepithema humile* (Lhu, Argentine ant), *Campanatus floridanus* (Cfl, Florida carpenter ant), *Pogonomyrmex barbatus* (Pba, red harvester ant), *Atta cephalotes* (Ace, Leafcutter ant), *Acromyrmex echinatior* (Aec, Panamian leafcutter ant), *Solenopsis invicta* (Sin, Red imported fire ant)*, Vollenhovia emery* (Vem, ant), *Ooceraea biroi* (Cbi, Clonal raider ant), *Wasmannia auropunctata* (Wau, Electric ant), *Polisted dominula* (Pdo, European paper wasp), *Neodiprion lecontei* (Nle, Red-headed pine sawfly), *Cephus cinctus* (Cci, Wheat stem sawfly); **Isoptera**: *Zootermopsis nevadensis* (Zne, Dampwood termite); **Lepidoptera:** *Bombyx mori* (Bmo, Silkmoth)*, Chilo suppressalis* (Csu, Asiatic rice borer), *Danaus plexippus* (Dpl, Monarch butterfly), *Heliconius melpomene* (Hme, Postman butterfly), *Plutella xylostella* (Pxy, Diamondback moth), *Manduca sexta* (Mse, Tobacco hornworm), *Cydia pomonella* (Cpo, Codling moth), *Vanessa tameamea* (Vta, Kamehameha butterfly), *Trichoplusia ni* (Tni, Cabbage looper), *Spodoptera litura* (Sli, Tobacco cutworm), *Galleria mellonella* (Gmo, Greater wax moth), **Odonata**: *Ischnura elegans* (Iel, damselfly), **Thysanoptera**: *Frankliniella occidentalis* (Foc, thrips). Capital letters above branches represent the amino acid residues at the ion channel selectivity filter (see main text and Fig. 3D).

#### 2.3. Supplementary Figure 3. qPCR analysis of RpTRPA5B

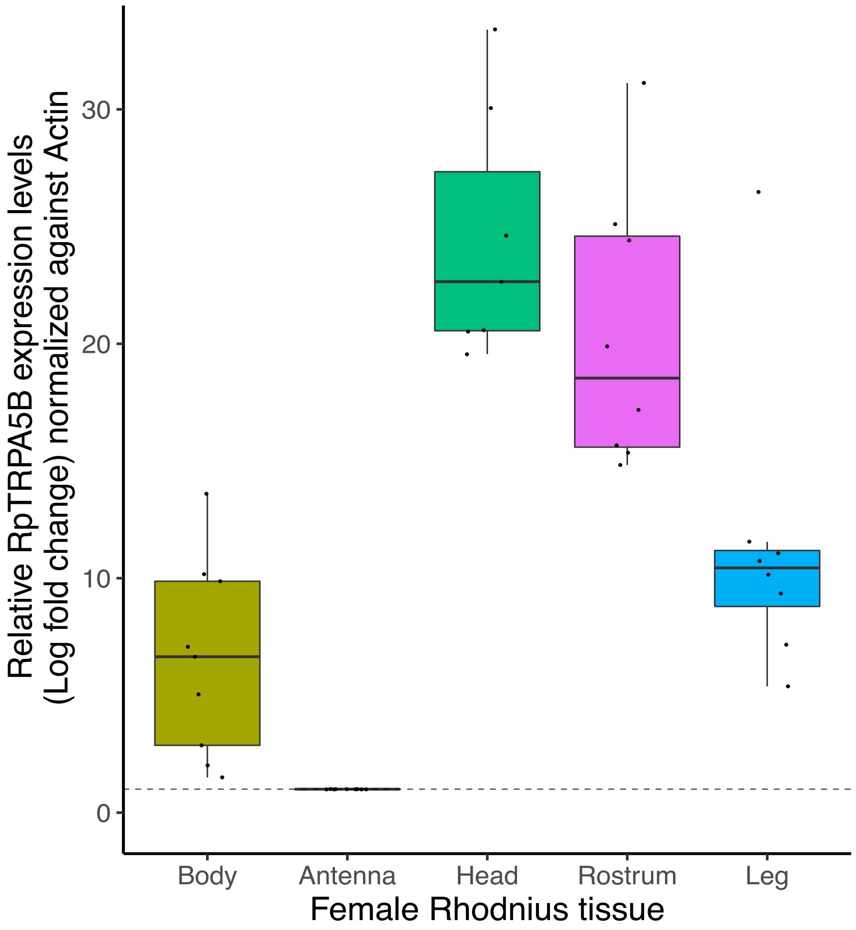


**Figure S3. qPCR analysis of RpTRPA5B.** $Log_2$ fold change expression was monitored by quantitative PCR and calculated from raw cycle threshold (Ct) values (±SEM, n= 9) (Supplementary Dataset, Table S3). Body tissue corresponds to Abdomen and Thorax minus legs. Box-and-whisker plots were visualized in RStudio 2021.09.2. Center horizontal bars show medians, box limit indicates upper and lower quartiles, whiskers show sample 1.5x interquartile range. Expression is calibrated using Antenna (log fold change =1) and normalized against Actin.

#### 2.4. Supplementary Figure S4: TRP whole cell and surface expression.

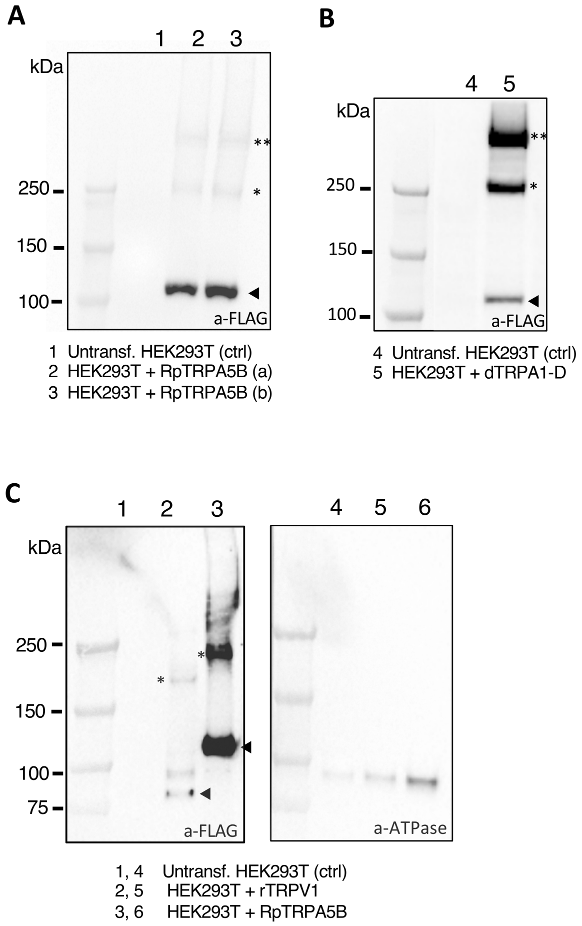


**Figure S4. TRP whole cell and surface expression. A-B**. RpTRPA5B anti-FLAG signals in whole cell lysates. Untransfected HEK293T cells (lanes 1,4) and HEK293T cells transfected with 2.5 µg pcDNA-FLAG-T2A-mRuby plasmid DNAs for (**A**) RpTRPA5B (a and b are two separate lysate samples, lanes 2,3), and (**B**) fruit fly dTRPA1-D (lane 5). Cells were collected 72 hrs after transfection. The protein ladder image taken from the same membrane is juxtaposed to the left of the immunoblot. One and two asterisks represent predicted dimeric and tetrameric TRP forms, respectively. The predicted monomeric molecular weight is indicated with a black arrowhead: RpTRPA5B, 127.78 kDa; fruit fly Dmel-TRPA1-D 138.82 kDa. **C.** Surface expression analysis of RpTRPA5B. Biotinylated surface protein eluates were run in parallel wells on the same SDS-PAGE gel, and after protein transfer, the membrane was split to probe TRP with a-FLAG (left) and ATPase with a-ATPase (right). Anti-FLAG levels in surface protein fraction are from non-transfected HEK293T cells (lane 1), cells expressing rTRPV1 (lane 2), and cells expressing RpTRPA5B (lane 3). Lanes 4 to 6 are the corresponding anti-ATPase biotin-surface fraction from non-transfected HEK293T cells (lane 4), cells expressing rTRPV1 (lane 5), and cells expressing RpTRPA5B (lane 6). One and two asterisks represent predicted dimeric and tetrameric TRP forms, respectively. The predicted monomeric weight is indicated with a black arrowhead (rTRPV1, 94.95 kDa; RpTRPA5B, 127.78 kDa).

#### 2.5. Supplementary Figure 5: Validation with rTRPV1 and dTRPA1-D

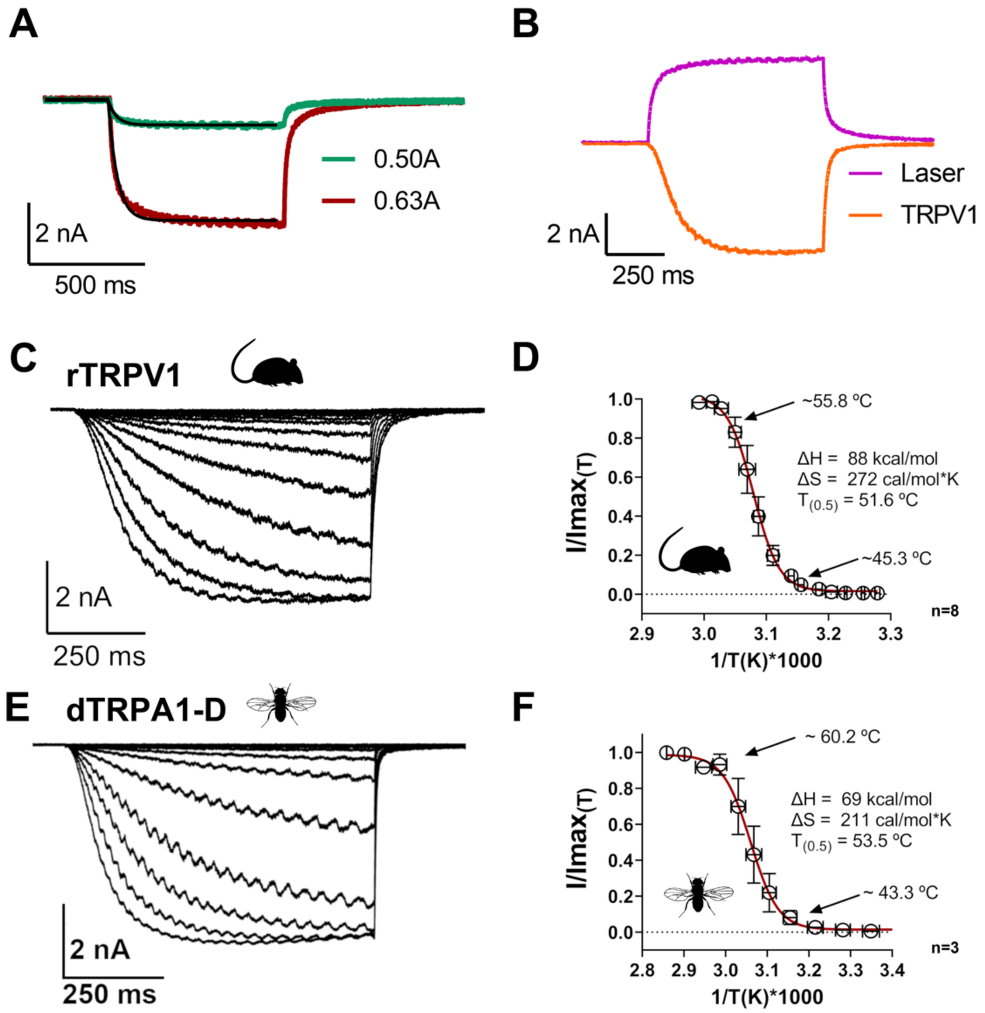


**Figure S5. Validation with rTRPV1 and dTRPA1-D**. **A.** Time course of the ionic current through the open pipette at -10 mV holding voltage when a current of 0.50A (green trace) and 0.63A (red trace) was applied to the laser diode. The temperature jumps for those currents correspond to 32.1 °C and 59.7 °C, from 22.6 °C, respectively. Both temperature jumps can be fitted by a mono-exponential function with a time constant of ~35 ms (black line). **B.** Time course comparison between the current through the open pipette (magenta trace) and a HEK293T cell expressing rTRPV1 channels under voltage clamp at -30 mV (orange trace), in response to a temperature jump from 22.6 °C to 59.7 °C (0.63A). The current through rTRPV1 channels is three times slower than the laser kinetics with a time constant of ~100 ms. The sinusoidal pattern observed within the current curves is inherent to the cyclic modulation of the laser's rapid 'on-off' cycles. **C.** Representative heat-activated current traces of a HEK293T cell expressing rTRPV1 under voltage clamp (holding voltage of -30 mV). The currents were elicited by temperature jumps from room temperature (22.6 °C) up to 59.7 °C with a duration of 700 ms. **D.** Fraction of rTRPV1 channels in the open state (Open probability, Po) as a function of the temperature. The Po vs 1/T was fitted to a Boltzman function (red line) with the midpoint of activation

($T_{0.5}$) reached at 51.6 °C. The van't Hoff plot estimates for rTRPV1 provides an activation enthalpy of the endothermic transition at 88.3 ± 9.4 kcal/mol and an entropic change associated to the temperature activation process at 271 ± 28 cal/mol*K. at -30 mV as a function of the temperature. These values in our expression cassette are very close to previously published values of ΔH = 85 kcal/mol, $T_{0.5}$ = 51 °C (Liu et al., 2003; Yao et al., 2010). These temperature values align with those reported in earlier studies for rTRPV1. The published threshold of activation for rTRPV1 is in the range 40-42 °C and corresponds to the temperature at which first observable currents are detected (Caterina et al., 1997; Tominaga et al., 1998; Liu et al., 2003). The channel emerging temperature recorded in our setup is at 40.9 °C when the first activation currents emerge over the dotted line (**D**). We calculated values of $T_{0.1}$ (i.e., the temperature for which there is a probability for 10% of channels to be open) at 45.3 °C (-30 mV), and $T_{0.5}$ at 51.6 °C (-30 mV), consistently with values reported in Yao et al 2010 ($T_{0.5}$ = 51°C, -60 mV). **E.** Representative heat-activated current traces of a HEK293 cell expressing dTRPA1-D channels under voltage clamp (holding voltage of -30 mV). The currents were elicited by temperature jumps from room temperature (19.4 °C) up to 63.5 °C with a duration of 700 ms. **F.** Fraction of dTRPA1-D channels in the open state (Open probability, Po) as a function of the temperature. The Po vs 1/T was fitted to a Boltzman function (red line) with the midpoint of activation ($T_{0.5}$) reached at 53.5 °C, the van't Hoff plot estimates for dTRPA1-D an activation enthalpy of the endothermic transition at 68.7 ± 13.1 kcal/mol and an entropic change associated to the temperature activation process at 211± 40 cal/mol·K at -30 mV. The stationary current at the end of the temperature pulse (last 100 ms) was used to calculate all the thermodynamics parameters. Data are presented as means ± standard deviation.

Caterina MJ, Schumacher MA, Tominaga M, Rosen TA, Levine JD. Julius D. 1997. The capsaicin receptor: a heat-activated ion channel in the pain pathway. Nature 389:816-824 (doi.org/10.1038/39807)

Liu B, Hui K, Qin F. 2003 Thermodynamics of Heat Activation of Single Capsaicin Ion Channels VR1 *Biophys J.* **85**(5): 2988–3006. (10.1016/S0006-3495(03)74719-5).

Tominaga M, Caterina MJ, Malmberg AB, Rosen TA, Gilbert H, Skinner K, et al. 1998. Neuron 21:531-543 (10.1016/s0896-6273(00)80564-4).

Yao J, Liu B, Qin F. 2010 Kinetic and energetic analysis of thermally activated TRPV1 channels. *Biophys J* **99**, 1743-1753. (10.1016/j.bpj.2010.07.022).

#### 2.6. Supplementary Figure S6: Open pipette calibration and fiber positioning

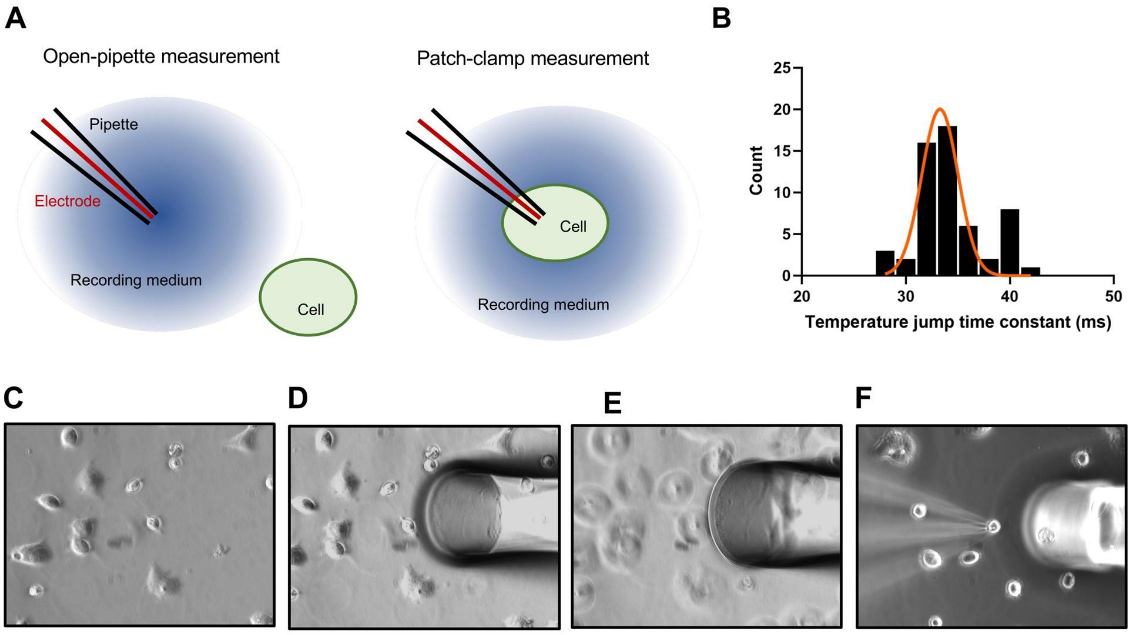


**Figure S6. Open pipette temperature calibration and optic fiber positioning. A.** Schematic representation of open pipette temperature calibration performed prior to each cell patch clamp experiment. First the open pipette is positioned under the center of the laser using the reference marks on the screen. Then we proceed to record the current through the open pipette elicited as a function of a series of near IR laser pulses. With these current traces we can estimate the temperature jumps magnitude that will be applied later to the target cell. After this the target cell is positioned in the center of the laser beam using the microscope stage translation (the optic fiber stays fixed). Once the cell is in the right position, we use the same pipette used for the calibration to carry out the electrophysiological recording, applying the same set of pulses used in the calibration **B.** Histogram of the time constants from the exponential fit to the open pipette current traces from the temperature jumps used in the rTRPV1 experiments (n=56), the mean time constant was 33.3 ± 1.8 ms. **C.** View of HEK293T cells seeded at low density on a glass cover in the recording chamber on the patch-clamp rig station. **D.** After finding a target cell co-expressing mRuby2 as a fluorescent marker, with the help of an automatized micromanipulator, the optic fiber is placed in the recording solution using the reference marks on the computer screen (not shown here) defined during the calibration with the visible laser. The depth in the solution is adjusted so that the fiber is directly above a single cell in the recording medium. **E.** The field of view is changed to align the patch clamp electrode to the reference marks. **F.** The target cell expressing mRuby2 is patched with the recording electrode, now visible on the left side, whereas the fiber is directly above the target cell. The relative position between the recording pipette and the fiber was established using a visible laser during the setup of the system, and is constant for all the experiments.

#### 2.7. Supplementary Figure 7: DALI Analysis

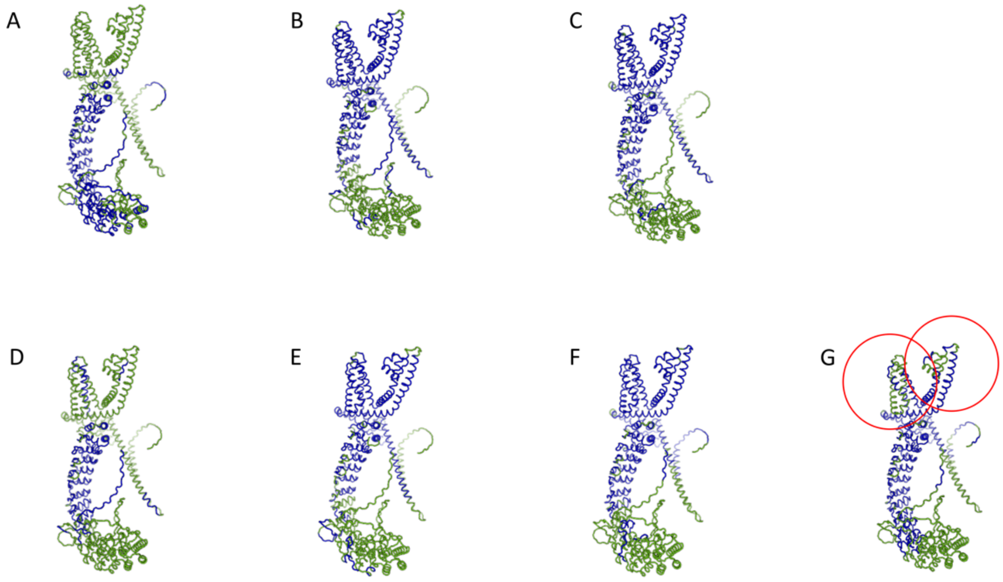


**Figure S7. DALI Analysis.** Pairwise structural alignment using the Dali server (http://ekhidna2.biocenter.helsinki.fi/dali/), of different monomers against *Rhodnius* TRPA5B. **A.** *Rhodnius* TRPA1, **B**. *Rhodnius* Painless, **C.** *Rhodnius* Waterwitch, **D.** *Drosophila* TRPA1, **E.** *Drosophila* Painless, **F.** *Drosophila* Waterwitch, **G.** *Drosophila* Pyrexia. The blue and green colors indicate regions with high and low structural similarity based on Z-values. The red circles in **G** indicate areas where the Pyrexia monomer stands out from Painless and Waterwitch, in that the pore region and the voltage-sensing like domain are less similar to TRPA5B. However, compared to Painless and Waterwitch the similarity in the ankyrin repeat domain (ARD) reaches further towards the N-terminus.

#### 2.8. Supplementary Figure S8: Validation of a tetrameric AlphaFold model of dTRPA1 with the experimentally determined structure dTRPA1 in state-2 (PDB ID 7YKS).

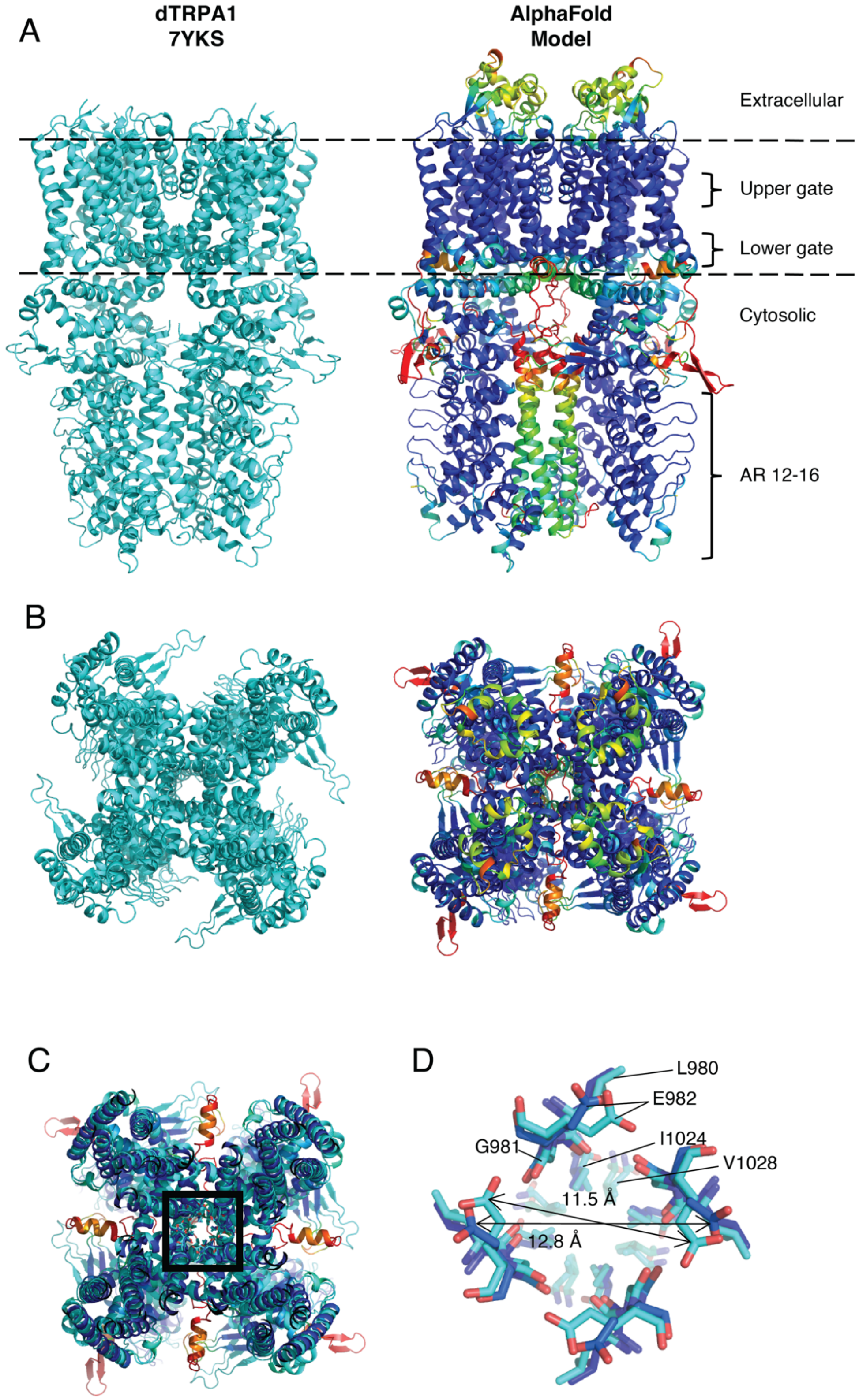

**Figure S8. Validation of a tetrameric AlphaFold model of dTRPA1 with the experimentally determined structure dTRPA1 in state-2 (PDB ID 7YKS)**. **A**. Side view of dTRPA1 with approximate boundaries of the transmembrane domain indicated by dashed lines. The N- and C-terminal regions that are not resolved in 7YKS were excluded in the prediction. Only the last five of the 17 ankyrin repeats (AR12-16) are visible in the structure and overall regions with low confidence in the model (red-yellow) are not resolved in the structure. **B.** Top view of **A. C.** Cross section of the pore region of the aligned structure and model in the same orientation as in B. Residues in the upper and lower gate regions are shown as sticks in the central box (numbering in 7YKS L980, G981, E982, and I1024, V1028, respectively). **D**. Close up of residues shown as sticks in **C**. The spatial location of these residues is very similar in the structure and the model. The main difference being the side chain of E982, which has a slightly more distal orientation relative to the center of the pore in the model. The distance between opposite Cδ of E982 is 12.8 Å, which is similar to the distances in dTRPA1 in state 2 (7YKS, 11.5 Å) and state-1 (7YKR, 12.7 Å). RMSD of residues in the upper and lower gate of the model with determined structures of dTRPA1 are provided in Table S6.

##### 2.9. Supplementary Figure S9: Distances between corresponding residues in the upper and lower gate.

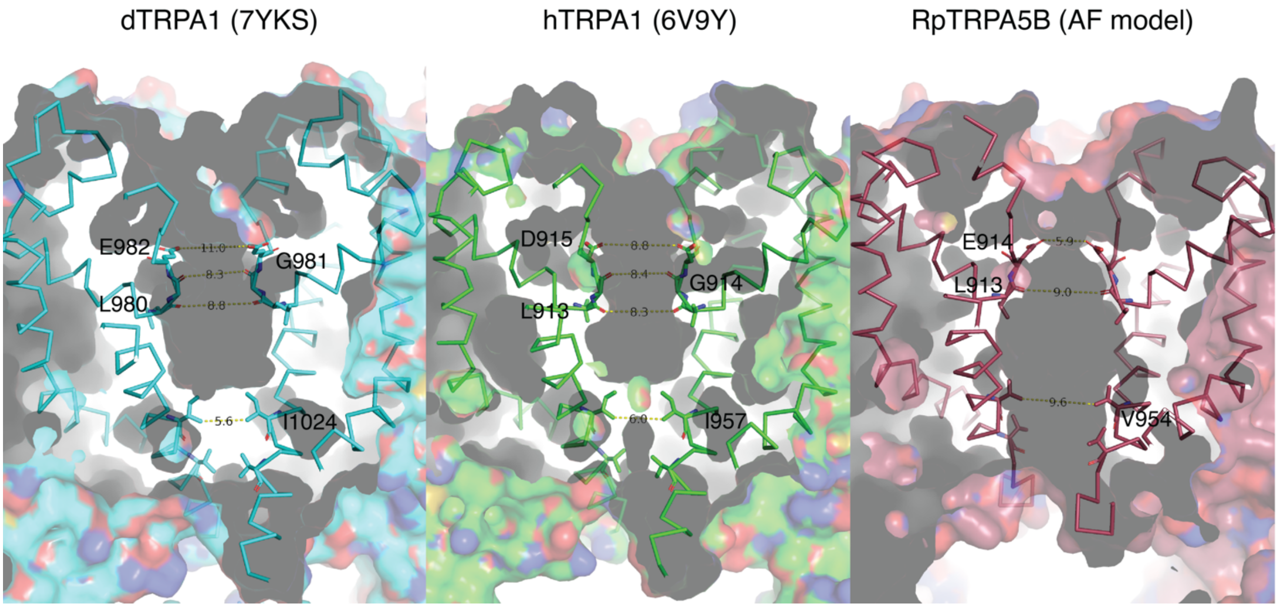


**Figure S9**. Distances between corresponding residues in the upper and lower gate in structures of TRPA1 from *Drosophila melanogaster* (dTRPA1; left) and *Homo sapiens* (hTRPA1; center) and the model of *Rhodnius prolixus*, RpTRPA5B (right).

##### 2.10. Supplementary Figure 10: Comparison of wild-type and predicted mutant RpTRPA5B

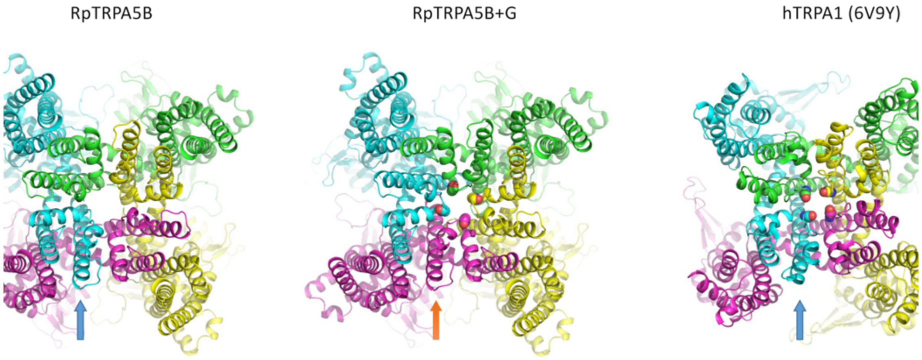


**Figure S10.** Extracellular view of truncated tetrameric AlphaFold model of RpTRPA5B (left), model of the corresponding mutated RpTRPA5B with the inserted glycine (spheres; middle) and hTRPA1 (6V9Y; corresponding glycine as spheres; right). With the extra glycine the overall fold of TRPA5B is predicted to be compromised and deviates from the generic fold of TRPs that is exemplified by hTRPA. In the mutant protein, helix S6 (orange arrow) is adopting the position of S6 (blue arrow) in a neighbouring monomer in the wild type. Thus, the loop between pore helices 1 and 2 forming the selectivity filter is predicted to be abnormally placed if adding the glycine without considering the potential impact of co-evolving sites. Cartoon representation coloured according to chain.
